# Reovirus recombination is highly selective, and its profiles are primarily dictated by viral gene segment identity

**DOI:** 10.1101/2025.10.23.684199

**Authors:** Alejandra Flores, Andrew Routh, Ryan D. Xavier, Kristen M. Ogden

**Affiliations:** Department of Pathology, Microbiology, and Immunology, Vanderbilt University Medical Center, Nashville, Tennessee, USA; ClickSeq Technologies LLC, Davis, California, USA; Department of Pediatrics, Vanderbilt University Medical Center, Nashville, Tennessee, USA

## Abstract

Recombination facilitates the generation of defective viral genomes (DVGs), truncated derivatives of the parental genome that require a helper virus to replicate. Recombination mechanisms are poorly understood for viruses with double-stranded RNA genomes. Two strains of the double-stranded RNA virus reovirus differ in the pattern of packaged DVGs during serial passage. To determine whether the polymerase complex or gene segment sequence contributes to these differences, we exchanged polymerase complexes between the two reovirus strains. We identified DVG patterns using RT-PCR and recombination junction profiles using ClickSeq. Reoviruses synthesized DVGs that maintained the 5′ and 3′ termini and contained large central deletions. The polymerase complex did not detectably affect DVG pattern following reovirus serial passage or viral recombination junction profiles. Instead, recombination junction profiles correlated with the identity of viral RNA gene segments, even in the presence of a non-native polymerase complex or virus background. Reovirus recombination junction start and stop sites often occur in regions of sequence microhomology. While we observed many instances of short stretches of identical nucleotides within a viral gene segment, only a select few positions were incorporated into recombination junctions. Overall, these data suggest that recombination events that can mediate reovirus DVG formation are highly selective, and properties of viral gene segments primarily dictate where recombination occurs. These observations suggest a model for double-stranded RNA virus recombination in which the polymerase pauses RNA synthesis and re-initiates further along the same template at a specific junction stop site that has sequence homology to the junction start site.

**IMPORTANCE:** Viral infection gives rise to defective viral genomes, which cannot complete a full replication cycle. Recombination facilitates defective viral genome generation but is understudied for RNA viruses with double-stranded genomes. We found that for a double-stranded RNA virus, reovirus, recombination occurs preferentially at specific sites in the genome that correspond with the identity of the gene segment. Recombination tends to initiate and terminate at sites sharing identical sequences. However, even if these short nucleotide sequences appear multiple times in a gene segment, only specific sites are used for recombination. Our results indicate that reovirus recombination is a highly orchestrated event in which individual gene segments contain the characteristics that drive recombination. These findings suggest that RNA properties, such as sequence and structure, drive recombination for double-stranded RNA viruses, likely through reinitiation after re-hybridization of the newly formed RNA product with the same RNA template molecule at a different location.

## INTRODUCTION

Most RNA viruses encode RNA polymerases that lack proofreading activities and can form defective viral genomes (DVGs). DVGs contain mutations, deletions, or changes in their genetic structure that prevent them from replicating without the assistance of a helper virus (1). DVGs can be packaged inside virions to form defective interfering particles (DIPs), which can interfere with viral replication. DIPs can reorient resources, outcompete wild-type viral replication, or activate the immune system (1–3). Viruses with relatively less DVG accumulation have been associated with more severe or fatal disease outcomes in humans and mice for negative-sense RNA virus influenza A virus, and a critical role for DVGs in modulating host interferon responses and symptom development has been suggested for positive-sense RNA virus SARS-CoV-2 (4, 5). Since they are replication-incompetent and can interfere with viral replication, DVGs and DIPs have been considered as potential therapeutics or vaccines for multiple viruses (6, 7). Although they have been well studied for positive-sense and negative-sense RNA viruses, relatively little is known about DVGs and DIPs of double-stranded RNA (dsRNA) viruses.

Mammalian orthoreovirus (reovirus) is a dsRNA virus that can package DVGs (8). Reovirus belongs to the order Reovirales and has garnered interest for its oncolytic therapy potential (9–11). Reovirus has a segmented dsRNA genome containing three large (L, 3.8-3.9 kb), three medium (M, 1.9-2.2 kb), and four small (S, 1.2-1.6 kb) segments (12). Reovirus gene segments contain an open reading frame (ORF) flanked by 5′ (12–32 nt) and 3′ (35-83 nt) untranslated regions (UTRs) (13). The ten reovirus dsRNA gene segments are present in equimolar proportions in virions, and particles tend to contain either all ten segments or none of them, indicating that packaging is highly orchestrated (14, 15). Numerous studies suggest the reovirus gene segment termini are important for packaging (16–22). Consistent with this observation, sequenced reovirus DVGs contain one or two large internal deletions but retain the 5′ and 3′ termini (8). However, the RT-PCR profiling used to obtain these sequences preferentially amplifies DVGs that conserve the termini and limits the detection of other types of DVGs. Two prototype human reovirus strains, T1L and T3D, differ in apoptosis induction capacity, viral factory morphology, and other characteristics (12). Serial passage of T1L and T3D in distinct evolutionary lineages in murine L929 fibroblasts (L cells) yielded viruses that packaged distinct subsets of DVGs, which we refer to as ‘patterns’ (8). While T1L had a consistent DVG pattern across evolutionary lineages, T3D packaged more diverse and changing subsets of DVGs. These observations suggest an inherent difference in the specificity with which T1L and T3D synthesize or package DVGs. Much work is needed to quantify the differences between T1L and T3D DVGs, elucidate the factors that contribute to these differences, and identify features of DVGs that promote their packaging.

For RNA viruses, recombination has been proposed as a major driver of DVG formation (2, 23, 24). For positive-sense RNA viruses and influenza viruses, deletion DVGs are detected that contain large internal deletions but often retain promoter regions and terminal sequences necessary for their replication and packaging (25). Deletion DVGs are thought to form when the viral RdRp dissociates from the template and reassociates either intra- or intermolecularly with the same or a homologous template (2, 23, 24). Copy-back DVGs, which are thought to occur when the viral polymerase dissociates from the template strand and reinitiates on the newly synthesized strand, are detected for non-segmented negative-sense RNA viruses (1, 25, 26). Recombination mechanisms for viruses containing segmented dsRNA genomes are largely undescribed. Inside reovirus particles, the RNA-dependent RNA polymerase (RdRp), λ3, forms a complex with its cofactor, µ2, which likely guides template RNA in and out of the RdRp, provides energy through its nucleoside triphosphatase activity, and acts as a helicase (27–31). Unlike simpler clamp-like RdRps, λ3 contains tunnels for RNA template and product entry and exit and cannot readily release and switch RNA templates (32–34). Following reovirus entry into cells, RNA transcription occurs inside the viral core, where the minus strands of genomic dsRNA segments are utilized as templates to synthesize positive-sense single-stranded RNA (+ssRNA) transcripts that are extruded into the cytoplasm. During particle assembly, the ten +ssRNA segments are used as templates for the synthesis of complementary minus-strand RNA to form genomic dsRNA (33). Like positive-sense RNA viruses, reovirus can form deletion DVGs (8). For rotavirus, another member of the order Reovirales, segment rearrangements have been associated with the RdRp; these rearrangements frequently involve partial head-to-tail RNA duplications that conserve the gene segment termini, and it is hypothesized that their synthesis involves RdRp dissociation and reassociation on the same template (35). Illumina RNA-seq analysis of reovirus strain T1L revealed recombination hot spots across the genome, and at least some junction sites appeared to occur near short regions of identical sequence, or microhomology (8). These observations suggested that reovirus recombination might occur at select sites. However, the studies were conducted for only two samples of a single reovirus strain. Furthermore, while unbiased, Illumina RNA-seq library preparation can introduce artifactual recombination events that account for a noteworthy portion of output sequence reads and influence computational analyses of DVGs (36). Thus, additional studies are needed to validate selective recombination events for dsRNA viruses like reovirus.

Since recombination events leading to DVG generation involve viral polymerase activity, we hypothesized that the observed differences between T1L and T3D in DVG patterns among evolutionary lineages (8) were due to differences in the viral polymerase complexes. Thus, we generated reoviruses in which the polymerase complex was exchanged between T1L and T3D. We exchanged segments L1 and M1, which respectively encode polymerase complex proteins λ3 and µ2, between T1L and T3D, rescued the recombinant reoviruses, and characterized them. These exchanges provided the opportunity to distinguish between the contributions of the RNA gene segments and the polymerase complex proteins they encoded to differences in DVG patterns. We serially passaged the viruses to determine DVG patterns in evolutionary lineages. We also sequenced packaged RNA in wild-type and polymerase complex-exchanged T1L and T3D reoviruses using an approach that minimizes artifactual recombination and analyzed the recombination junction sites. We found that TIL and T3D differ in their recombination junction site preferences, but the phenotype segregates genetically with the identity of the gene segment rather than the polymerase complex. At recombination junction start and stop sites, we detected conservation of positional identity but not of a specific sequence motif directing recombination. Although sequences exhibiting microhomology were found to appear multiple times in a gene segment, preferred sites were used for recombination. Together, these observations suggest that RNA primary sequence is required for DVG synthesis and selection.

## RESULTS

### Polymerase complex-exchanged reoviruses exhibit distinct replication phenotypes

To investigate how the polymerase complex influences packaged DVG populations, we engineered T1L and T3D reoviruses with exchanged polymerase complexes. We exchanged the L1 and M1 gene segments of reovirus strains T1L and T3D using reverse genetics to generate T1L-T3pol and T3D-T1pol (**Fig. 1A**) (37, 38). To confirm that both L1 and M1 segments were swapped between the strains, we utilized electropherotyping to separate viral gene segments based on size and electrical charge (**Fig. 1B**). For T1L-T3pol, gene segments L1 and M1 migrated more slowly in the gel compared to the T1L gene segments, instead resembling those of T3D. In contrast, for T3D-T1pol, gene segments L1 and M1 migrated faster and resembled those of T1L. These results indicate that polymerase complex-exchanged reoviruses were rescued using reverse genetics.

**Figure 1.**
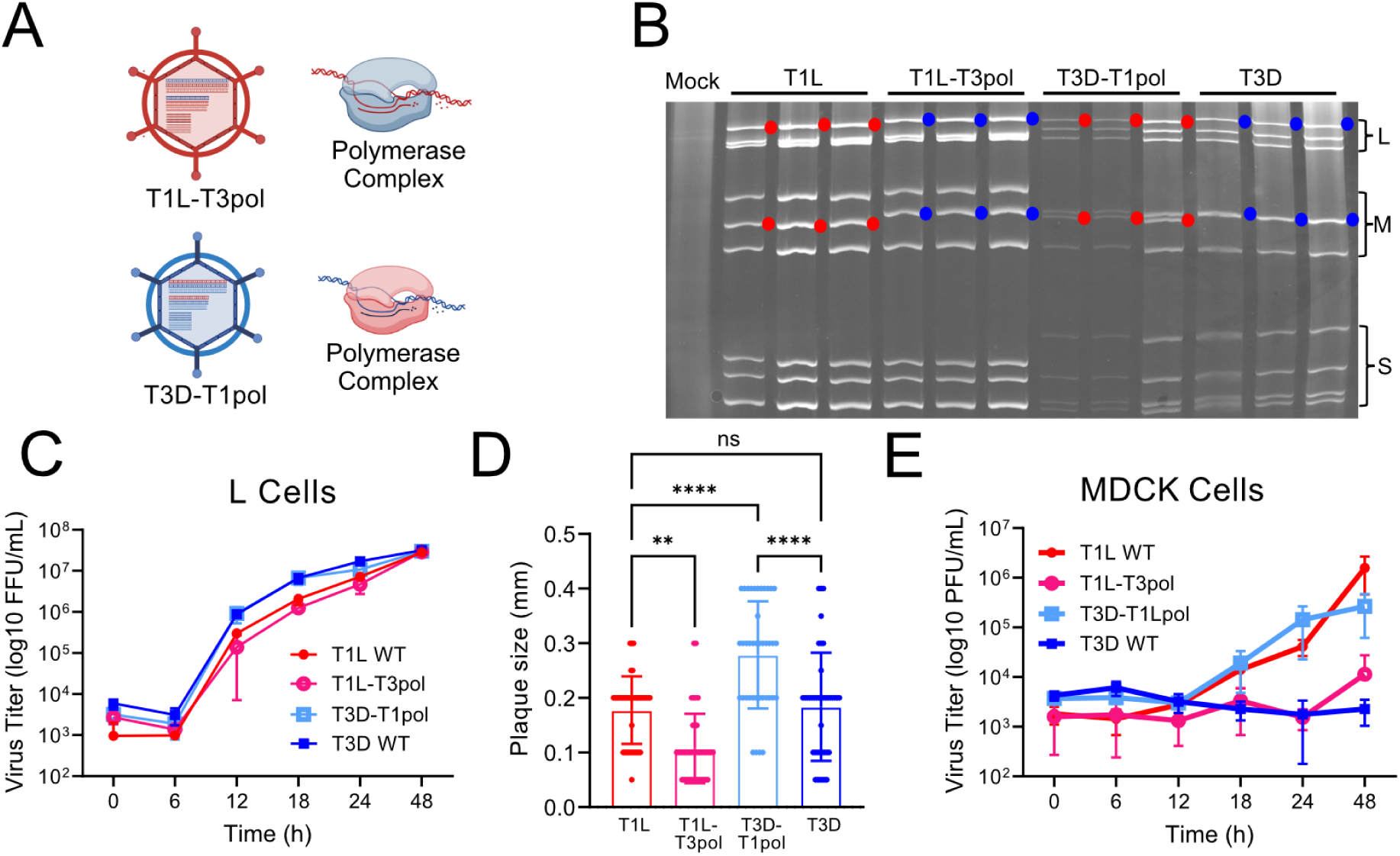
Polymerase complex-exchanged reovirus characterization. (A) Schematic of polymerase-exchanged viruses and their polymerase complexes. The origin of viral gene segments, represented as parallel lines inside the particle, the virus background, represented by the particle, and polymerase complex are indicated by color (red, T1L; blue, T3D). Created using Biorender.com (B) Electropherotypes of T1L, T3D, and polymerase-exchanged reoviruses. RNA extracted from cells infected with the indicated virus was resolved by SDS-PAGE. Blue circles indicate T3D L1 or M1 gene segments. Red circles indicate T1L L1 or M1 gene segments. (C) Replication kinetics of parental and polymerase-exchanged reoviruses in L cells. Viral titers were determined by fluorescent focus assay. Error bars = SD, n = 3 clones per virus. Not statistically significant by two-way ANOVA. (D) Plaque size of parental and polymerase-exchanged reoviruses in L cells. ns, not significant; **, P < 0.01; ****, P < 0.0001 by one-way ANOVA. (E) Replication kinetics of parental and polymerase-exchanged reoviruses in MDCK cells. Viral titers were determined by plaque assay. Error bars = SD, n = 3 clones per virus. Not statistically significant by two-way ANOVA.

We analyzed the replication characteristics of polymerase-exchanged reoviruses using two different cell lines. In previous studies, T1L and T3D replicated with similar efficiency in L cells, but T1L replicated more efficiently in canine epithelial MDCK cells compared to T3D; these differences in growth segregated with the viral L1 and M1 segments (39, 40). To determine the effects of polymerase complex exchange on viral replication, we adsorbed L cells or MDCK cells with three clones each of T1L, T3D, T1L-T3pol, or T3D-T1pol at an MOI of 0.1 PFU/cell, washed, and incubated. Then, we quantified viral titers in total cellular lysate. In L cells, T1L-T3pol had similar replication kinetics to T1L, and T3D-T1pol had similar replication kinetics to T3D (**Fig. 1C**). However, spread of the four viruses differed in a plaque assay, in which viruses undergo multiple rounds of replication. The plaque size of T1L-T3pol was smaller than that of T1L, while T3D-T1pol had the largest plaque size among the four viruses, suggesting that the polymerase complex can confer a multi-cycle replication disadvantage or advantage in its non-native context (**Fig. 1D**). Consistent with previous results, T1L and T3D-T1pol were capable of replicating in MDCK cells, while T3D and T1L-T3pol largely failed to replicate (**Fig. 1E**) (39, 40). These results confirm the identities of reoviruses with exchanged polymerase complexes and suggest that their replication phenotypes can be influenced by the polymerase complex.

### Exchanging the polymerase complex fails to exchange lineage-specific DVG segment patterns

We previously found that DVG patterns differed for serially-passaged T1L and T3D reoviruses, with T1L accumulating similar DVGs and T3D accumulating more diverse DVGs in distinct evolutionary lineages (8). To determine whether the polymerase complex contributes to lineage-specific differences in packaged DVG patterns, we serially passaged T1L, T3D, and the polymerase complex-exchanged reoviruses in L cells. We initiated the passage series by adsorbing L cells at an MOI of 0.3 PFU/cell in triplicate to compare independent evolutionary lineages. For passage 1 (P1), the inoculum was recovered using reverse genetics and expanded once in L cells to minimize the presence of DVGs and generate sufficient virus for the infections. After adsorption, we incubated the cells for 48 h prior to lysis. Then, we infected a new round of cells with a fixed volume of the lysate to allow a high MOI for DVG formation. We repeated this procedure for 10 serial passages (**Fig. S1A**). We also conducted mock infections in triplicate, using PBS for the inoculum, to use as controls. We determined viral titers for each passage and lineage. Each analyzed virus showed distinct titer trends (**Fig. S1B-E**). In most cases, titers of the three independent lineages of a given virus exhibited similar increases or decreases across the passages.

To detect DVGs generated during serial passage and their patterns, we used an RT-PCR approach, in which RNA molecules containing the 5′ and 3′ termini of a specific viral gene segment were amplified and visualized for individual serial passages and lineages. The assay conditions permitted detection of small RNA molecules that preserved the segment termini, which have previously been shown to include DVGs (8). The selected segments vary in size and represent both exchanged and non-exchanged segments. Using this approach, full-length reovirus S and M segments, which are < 2.5 kb in length, were consistently detected, although the longer L segments were less consistently detected (**Figs. 2, S2-S4**). To determine whether our primers amplify host RNA, we conducted RT-PCR reactions using RNA extracted from the three mock-infected passages and for each set of primers. We found no evidence of off-target amplification for any set of primers (**Fig. S5**). For segment L1, a DVG of ∼ 300 bp was identified for all passages and lineages (**Fig. 2**). For T1L and T1L-T3pol, the ∼ 300-bp DVG was predominant (**Fig. 2A-B**). For T3D-T1pol and T3D, additional L1 DVGs of varying size were detected in specific passages and lineages (**Fig. 2C-D**). Thus, as previously observed, for the L1 segment, the DVGs detected were relatively consistent among T1L lineages and more variable among T3D lineages (8). However, L1 DVG variability does not appear to have been exchanged with the polymerase complex, as the T1L-T3pol lineages remained consistent, with a few additional DVGs detected in P9 and P10 of T1L-T3pol lin3, and the T3D-T1pol lineages were more variable (**Fig. 2**). For segment L3, most DVGs ranged from ∼ 100 bp to 800 bp in length, and some DVGs became more prominent in later passages (**Fig.S2)**. For each passaged reovirus, one to few L3 DVGs of similar size were detected across most passages for the three lineages of a virus, while the others varied in size, although L3 DVG sizes were more consistent across lineages for T1L. For segment M1, one or a small number of DVGs of the same size were detected in multiple passages for all lineages of a given virus (**Fig. S3**). For segment S4, a DVG of ∼ 150 bp was consistently detected for each passage and lineage of all four viruses (**Fig. S4**). For segments M1 and S4, products higher in size than the full-length segment were occasionally detected, which might have arisen from duplication events (**Figs. S3-S4**). These studies support the previous observation that DVG patterns revealed by RT-PCR generally tend to be less variable among lineages for T1L than T3D, at least for segments L1 and L3 (**Figs. 2 and S2**) (8). All the viruses in the study formed DVGs that can be detected by RT-PCR. However, the DVG patterns observed for T1L-T3pol and T3D-T1pol fail to conclusively support a role for the polymerase complex in the variety of DVGs synthesized and packaged during serial virus passage.

**Figure 2.**
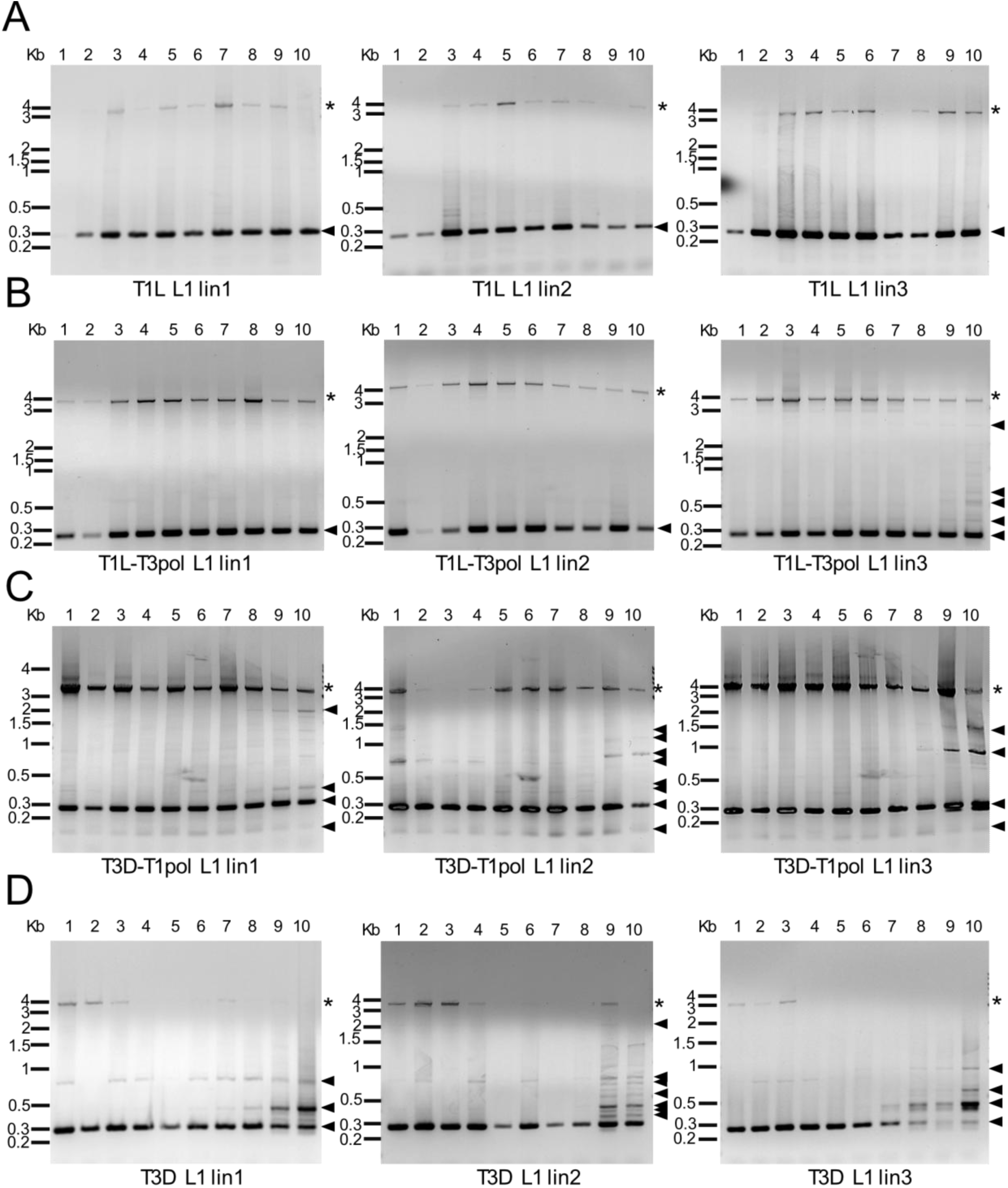
Serial passage lineage 1 to 3 reovirus gene segment L1 patterns. RNA extracted from serial passages was used as a template for RT-PCR reactions in which primers targeting terminal sequences in the L1 segments of both T1L and T3D were included. Products from T1L (A), T1L-T3pol (B), T3D-T1pol (C), or T3D (D) P1-P10 were resolved in 1.2% ethidium bromide-stained agarose gels. The evolutionary lineage is indicated. Black asterisks indicate the position of the full-length segment. Triangles indicate products smaller than the full-length segment.

### DVGs from polymerase-exchanged viruses contain large internal deletions

Our previous study determined that reovirus DVGs from segments L1, M1, and S4 contain relatively short intact termini and one or two large internal deletions (8). Resolved RT-PCR products from the current study revealed that reoviruses package DVGs derived from segments L1 and S4 that persist across the passage series (**Figs. 2, S4**). To determine whether the persistent DVGs synthesized and packaged by different viruses are identical, we extracted and sequenced RT-PCR-amplified products from segments L1 and S4 using the Sanger method for polymerase-exchanged reoviruses. Analysis of the sequences identified DVGs with internal deletions (**Fig. 3**). These DVGs are 276-282 bp (L1) and 147 bp (S4) in total length. The L1 DVGs each contain an internal deletion of 3579 bp in slightly different positions in terms of nucleotide number but at an identical junction in terms of sequence. The L1 5′ UTR is completely conserved along with 242 nt of the ORF and the last 16 nt of the 32-nt 3′ UTR, ∼ 7% of the full-length segment. The sequenced T3D-T1pol L1 DVG was identical to a T1L L1 DVG sequenced in our previous study (8). The S4 DVGs from T1L-T3pol and T3D-T1pol contained an identical internal 1050 bp deletion. The S4 5′ UTR is completely conserved along with 101 nt of the ORF and the last 14 nt of the 66-nt 3′ UTR, ∼ 12% of the full-length segment. These observations confirm that reovirus DVGs can contain large internal deletions and suggest that while both termini may be important for packaging, only a portion of the 3′ UTR is required. Detecting DVGs that are identical across passages, lineages, and virus backgrounds also suggests that DVGs may be selectively synthesized or packaged.

**Figure 3.**
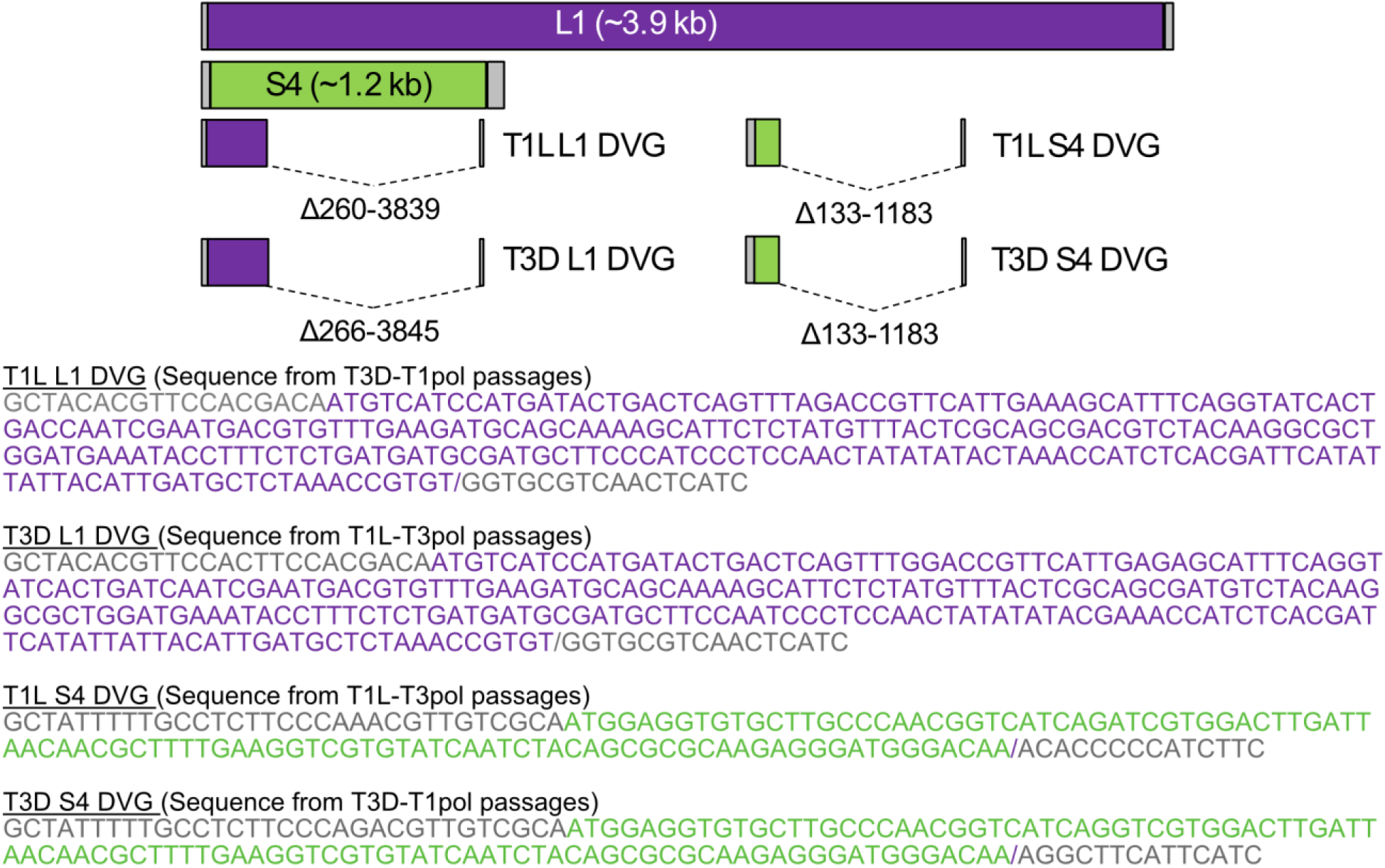
Sequences of isolated DVGs from serial passages. Schematics, drawn to scale, of full-length reovirus L1 and S4 segments and of sequenced T1L-T3pol and T3D-T1pol reovirus DVGs. The ORF is colored purple (L1) or green (S4). The 5′ and 3′ UTRs are colored gray. Central deletions were identified for each DVG. Dotted lines represent the missing sequences. Below the schematics are the DVG sequences, with DVG junctions indicated by “/”.

### Exchanging the polymerase complex fails to exchange reovirus recombination profiles

Although informative, DVG profiling using RT-PCR is biased by primers that bind reovirus gene segment termini and fails to provide sequence information. This approach can also selectively amplify small products relative to longer ones, misrepresenting their relative quantities. Thus, to elucidate sequence features of packaged DVGs and contributions of the polymerase complex to DVG synthesis and/or packaging, we deep-sequenced RNA encapsidated within T1L, T1L-T3pol, T3D-T1pol, and T3D reovirus particles using ClickSeq (36). ClickSeq is an RNAseq approach that uses bioconjugation as an alternative to fragmentation and enzymatic adaptor ligation during library preparation. This process reduces artifactual recombination to ∼2.5 events per million reads, even without careful optimization of PCR conditions (36). Thus, identified recombination junctions are likely to represent biologically relevant RNA products. Although ClickSeq has many benefits, a caveat is that the 3’-azido nucleotide (AzNTP):dNTP ratio used in library preparation yields cDNA fragments ∼ 300-500 nt in length (36). Thus, the capacity to detect the smaller products that are readily amplified using RT-PCR, including small DVGs (**Figs. 2 and S2-S4**), is limited.

Following multiple rounds of replication in L cells, we independently purified reovirus particles from three clones of each virus and treated them with benzonase to remove non-encapsidated RNA. Then, we extracted encapsidated RNA, prepared ClickSeq libraries from each RNA sample, and sequenced the products on an Illumina NovaSeq platform. Sequence coverage depth was high for every virus preparation and segment (**Fig. S6** and data not shown). Coverage depth varied positionally by ∼10-100-fold, with variation highly similar among replicates. Coverage biases may result from differences in the efficiency of random priming, reverse transcription, and PCR amplification and may be influenced by sequence-dependent factors, such as GC-content or RNA structure. Overall coverage was within normal limits, and the profiles generated were even. Greater than 99.9% of reads mapped to viral gene segments, demonstrating that reovirus virions had been sufficiently purified (**Table 1**). We used a viral recombination mapping algorithm (*ViReMa*) to identify recombination junction sites in the sequencing data, which were defined as deletions or insertions greater than five bp in length flanked both upstream and downstream by a minimum 25-bp high-quality alignment (**Fig. 4A**) (41, 42). We detected an average viral genome-wide recombination frequency (JFreq = reads with mapped junctions per 10,000 mapped reads) of 1.02-2.15, which is substantially lower than reported for betacoronaviruses (43) for example, and many noncanonical recombination junctions (leading to non-full-length products) in individual viral gene segments were identified for each virus (**Table 1**). Although the full-length sequences of DVGs cannot be determined from these data due to the short reads inherent to the Illumina sequencing platform, in at least one replicate for each of the four viruses, we detected the L1 recombination start site shown in **Fig. 3**, and for T3D, we detected the L1 recombination stop site. For T1L-T3pol, we detected the S4 recombination start site that is shown in **Fig. 3** with a stop site at nucleotide position 1146. Thus, some of the same recombination sites we found in sequenced DVGs were identified using *ViReMa*. These recombination sites were not identified in all samples, likely due to the reduced capacity of ClickSeq to capture very small products (36), including the smaller DVGs efficiently amplified by RT-PCR (**Figs. 2 and S2-S4**).

**Figure 4.**
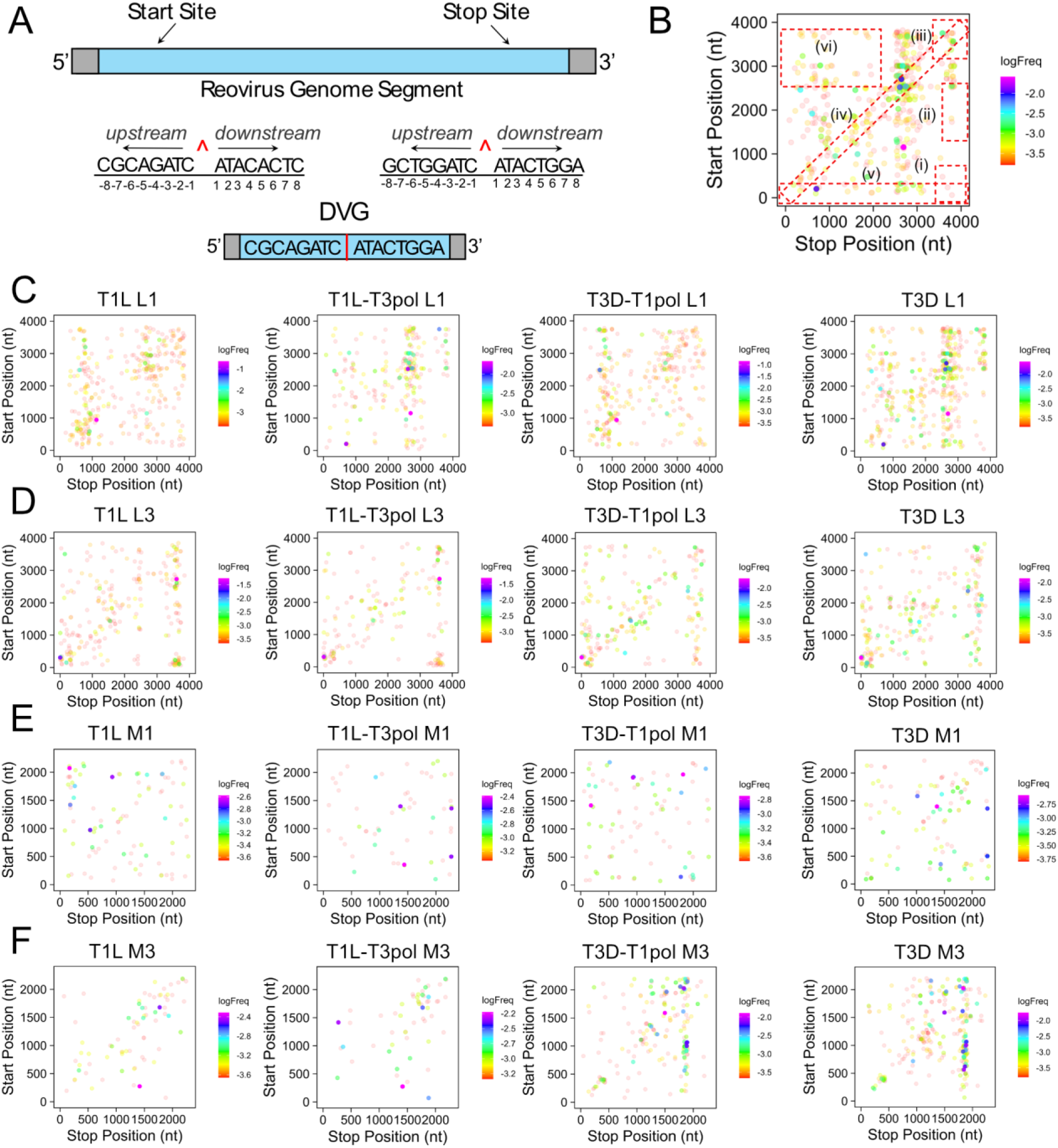
Reovirus recombination junction site location and frequency. (A) Schematic of a recombination junction site in the context of the parental genome segment, in which the 5’ and 3’ termini are conserved (bottom). The junction is labeled with a red caret (^), and example sequences are shown. Sequences upstream of the junction start site and downstream of the junction stop site may form a final product that is missing the internal genome sequence (top). (B) Schematic of junction scatter plots indicating the different cluster within the plot marked by dashed boxes: (i) 5’ → 3’ (ii) mid-genome → 3’ (iii) 3’ → 3’ (iv) local deletions/microindels (v) 5’ UTR → rest of genome (vi) duplications. (C-F) Recombination junction site location and frequency in sequenced T1L, T1L-T3pol, T3D-T1pol, and T3D virion RNA for gene segments L1 (C), L3 (D), M1 (E), and M3 (F). Junction sites are indicated by dots whose position corresponds to upstream and downstream sequences that are merged to form a novel junction. Junction frequency is indicated by dot color, according to the legend at the right of each image. Results from a representative clone for each virus are shown.

**Table 1.** Alignment statistics.

| Alignment statistic | Segment | T3D 1 | T3D 2 | T3D 3 | T3D-T1pol 1 | T3D-T1pol 2 | T3D-T1pol 3 | T1L-T3pol 1 | T1L-T3pol 2 | T1L-T3pol 3 | T1L 1 | T1L 2 | T1L 3 |
| --- | --- | --- | --- | --- | --- | --- | --- | --- | --- | --- | --- | --- | --- |
| % reads mapped to virus |  | >99.9% | >99.9% | >99.9% | >99.9% | >99.9% | >99.9% | >99.9% | >99.9% | >99.9% | >99.9% | >99.9% | >99.9% |
| Total reads |  | 25260410 | 28604063 | 30095851 | 26658380 | 27516061 | 29361973 | 30308219 | 21458634 | 25077376 | 26946402 | 29957691 | 28476911 |
| Jfreq <sup>a</sup> |  | 1.98 | 2.15 | 1.24 | 1.82 | 1.64 | 1.91 | 1.41 | 1.02 | 1.28 | 1.42 | 1.51 | 1.05 |
| # reads with recombination events |  |  |  |  |  |  |  |  |  |  |  |  |  |
|  | L1 | 1158 | 1513 | 1321 | 1390 | 1531 | 1802 | 1164 | 583 | 1018 | 973 | 1885 | 1081 |
|  | L2 | 1197 | 1424 | 591 | 1016 | 1026 | 1166 | 578 | 212 | 249 | 343 | 414 | 242 |
|  | L3 | 599 | 749 | 681 | 752 | 595 | 743 | 926 | 499 | 412 | 542 | 848 | 667 |
|  | M1 | 184 | 237 | 140 | 188 | 159 | 252 | 210 | 108 | 139 | 207 | 184 | 94 |
|  | M2 | 190 | 234 | 180 | 189 | 126 | 182 | 390 | 202 | 302 | 503 | 368 | 258 |
|  | M3 | 1043 | 1385 | 441 | 769 | 720 | 1009 | 207 | 120 | 181 | 270 | 158 | 115 |
|  | S1 | 262 | 135 | 47 | 164 | 49 | 76 | 310 | 214 | 581 | 707 | 322 | 234 |
|  | S2 | 107 | 137 | 69 | 90 | 63 | 80 | 138 | 61 | 95 | 96 | 86 | 59 |
|  | S3 | 178 | 210 | 202 | 154 | 170 | 177 | 75 | 67 | 142 | 110 | 67 | 63 |
|  | S4 | 94 | 129 | 71 | 139 | 83 | 118 | 284 | 122 | 102 | 67 | 200 | 183 |
| # reads mapped to viral gene segment |  |  |  |  |  |  |  |  |  |  |  |  |  |
|  | L1 | 5,106,848 | 5,824,267 | 7,563,379 | 5,830,947 | 6,058,222 | 6,264,212 | 5,801,679 | 3,638,187 | 3,704,550 | 4,417,355 | 5,891,865 | 5,463,064 |
|  | L2 | 6,662,826 | 6,978,216 | 7,265,286 | 6,848,569 | 7,341,564 | 7,322,106 | 4,720,881 | 2,995,701 | 2,889,015 | 3,371,773 | 4,430,912 | 4,339,775 |
|  | L3 | 3,162,471 | 3,729,886 | 4,220,554 | 3,152,933 | 3,516,584 | 3,631,513 | 4,330,594 | 3,031,626 | 2,864,871 | 3,379,424 | 4,377,796 | 4,097,063 |
|  | M1 | 2,010,161 | 2,401,261 | 2,171,824 | 2,366,934 | 2,368,327 | 2,690,442 | 2,186,178 | 1,630,315 | 1,863,444 | 2,278,138 | 2,344,993 | 2,167,978 |
|  | M2 | 1,988,216 | 2,285,659 | 2,066,003 | 1,943,737 | 1,936,186 | 2,209,843 | 5,631,920 | 3,996,113 | 3,903,288 | 4,333,452 | 5,549,127 | 5,224,089 |
|  | M3 | 2,331,392 | 2,867,077 | 2,901,518 | 2,395,419 | 2,473,377 | 2,921,266 | 1,715,465 | 1,352,975 | 1,606,108 | 1,774,066 | 1,645,633 | 1,639,310 |
|  | S1 | 1,321,199 | 1,452,403 | 1,090,694 | 1,336,338 | 1,258,269 | 1,428,736 | 2,284,308 | 2,002,678 | 3,496,513 | 3,331,144 | 2,410,284 | 2,278,743 |
|  | S2 | 1,059,981 | 1,048,569 | 824,693 | 1,122,301 | 956,053 | 1,121,603 | 1,550,675 | 1,161,403 | 1,783,005 | 1,500,017 | 1,435,369 | 1,241,740 |
|  | S3 | 500,594 | 590,814 | 767,147 | 549,583 | 524,739 | 570,384 | 816,523 | 675,750 | 1,445,408 | 1,328,825 | 799,117 | 921,942 |
|  | S4 | 1,112,954 | 1,342,805 | 1,219,969 | 1,099,702 | 1,079,295 | 1,198,661 | 1,266,340 | 971,054 | 1,519,289 | 1,230,540 | 1,069,488 | 1,100,512 |

<sup>a</sup> Number of reads containing junctions per 10,000 mapped reads

To visualize the locations and frequencies of recombination junctions for individual segments, we generated junction scatter plots (**Figs. 4 and S7-S10**). These plots revealed noncanonical junctions within each viral gene segment that suggest the presence of DVGs containing deletions or duplications (**Fig. 4B**), which have previously been detected for dsRNA viruses (8, 44–47). For segment L1, the identified junction profiles were visually different between the T1L and T3D strains (**Figs. 4C, S7**). We use the term “profile” to refer to the distribution of recombination junction identities for a given viral gene segment and their frequencies. The T1L-T3pol junction profiles for segment L1 partially resembled those of T3D, with which it shares a T3D L1 segment, while T3D-T1pol junction profiles for segment L1 partially resembled those of T1L, with which it shares a T1L L1 segment. For segment L3, the recombination profiles between T1L and T3D also visually differed (**Figs. 4D, S8**). However, the T1L-T3pol L3 junction profiles visually resembled those of T1L, with which it shares the T1L L3 segment. The T3D-T1pol junction profiles visually resembled those of T3D, which both contain the T3D L3 segment. For segment M1, visual similarities among junction profiles were not as evident compared to other segments (**Figs. 4E, S9**). For segment M3, the junction profiles resembled those of L3, with those of T1L and T1L-T3pol or T3D and T3D-T1pol more visually similar to one another (**Figs. 4F, S10**). Overall, viruses that share the same gene segment presented visually similar recombination profiles for that given segment, even in the presence of a different polymerase complex. Furthermore, the junction profiles were retained when segments were encapsidated in a different virus background, indicating that some property of the RNA segment dictates the recombination junction profile.

Previous research in our lab suggested that T1L recombination might occur preferentially at distinct sites, but only two samples were investigated for a single virus strain (8). To determine whether the genome contains ‘recombination hotspots’, we calculated the recombination frequency for segments L1, L3, M1, and M3 for three clones of each of our four viruses. The frequency of the start or stop position of each junction was evaluated by dividing the frequency of reads mapping to a recombination junction by the total mapped reads. The total reads were calculated as the sum of the depth. Results are shown as the mean of the three clones (**Figs. 5 and S11**) to facilitate comparison between the viruses and segments, and they are shown individually (**Figs. S12-S19**) to compare reproducibility among clones. Individual peaks representing regions of high frequency were not identical in all clones, but distinguishing recombination frequencies were typical for each virus. For segment L1, recombination start and stop sites occurring with the highest frequencies for T1L tended to be located at identical places in the gene segment as those of T3D-T1pol, whereas recombination start and stop sites occurring with the highest frequencies for T3D tended to be located at identical places in the gene segment as those of T1L-T3pol (**Figs. 5A, S11A, and S12-S13**). In contrast, for segment L3, recombination start and stop sites occurring with the highest frequencies for T1L tended to be located at identical places in the gene segment as those of T1L-T3pol, while recombination start and stop sites occurring with the highest frequencies for T3D tended to be located at identical places in the gene segment as those of T3D-T1pol (**Figs. 5B, S11B, and S14-S15**). For segment M1 start and stop site frequencies across the segment, T1L appeared most visually similar with T3D-T1pol, whereas T3D resembled T1L-T3pol (**Figs. 5C, S11C, and S16-S17**). Finally, for segment M3 start and stop sites, T1L and T1L-T3pol both tended to share positions of high frequency recombination sites, while T3D and T3D-T1pol shared such positions (**Figs. 5D, S11D, and S18-S19**). In general, like the recombination junction patterns (**Figs. 4 and S7-S10**), the recombination frequency peak patterns of T1L-T3pol and T3D-T1pol visually resembled those of the strain of origin for each given segment investigated, regardless of whether the segment was associated with a different polymerase complex or a new virus background. Though qualitative, these observations suggest that reovirus recombination profiles segregate with the reovirus gene segment identity independent of the source of the polymerase or other viral proteins.

**Figure 5.**
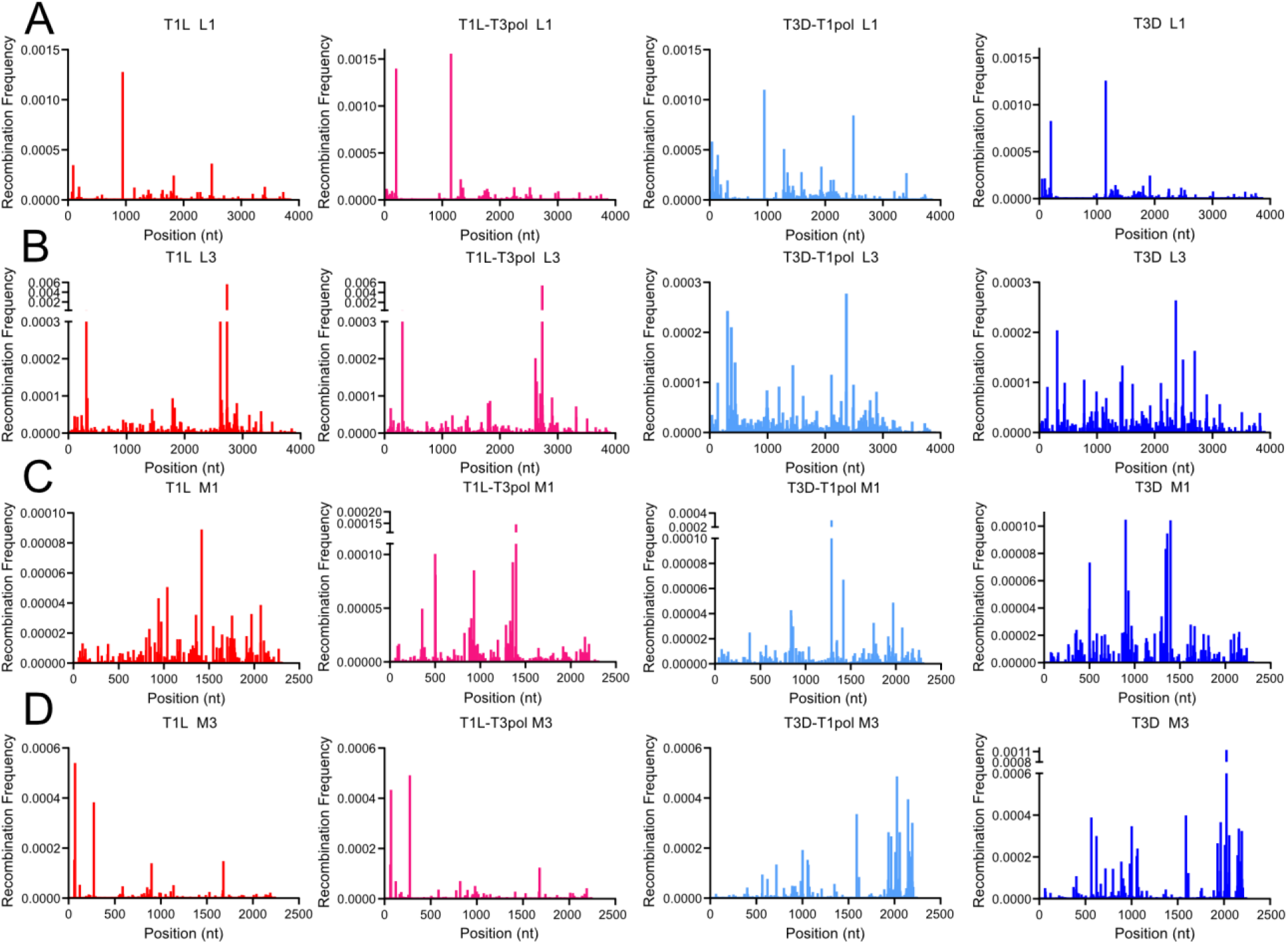
Reovirus recombination start-site frequency across gene segments. Recombination start-site frequency at each nucleotide position across L1 (A), L3 (B), M1 (C), and M3 (D) segments from T1L, T1L-T3pol, T3D-T1pol, and T3D. Mean positional recombination frequency for sequenced virion RNA from three clones per virus is shown.

### Principal component analysis reveals that recombination profiles segregate with the reovirus gene segment independent of the polymerase complex

Recombination frequencies and junction scatter plot data visually suggest similar profiles between reoviruses containing segments derived from the same virus regardless of the origin of other viral proteins. To quantitatively assess similarities in junction profiles among the four viruses, we conducted a principal component analysis (PCA) for segments L1, L3, M1, and M3. For each virus and replicate, we analyzed the collection of unique recombination junctions and their frequencies. The loadings table was utilized to compare the similarity of viral recombination junction profiles for each given segment. For segment L1, T1L clustered tightly with T3D-T1pol, T3D clustered tightly with T1L-T3pol, and the proportion of variance for the two first principal components was 92.93, indicating that nearly all variance in the dataset is explained by the first two components (**Fig. 6A**). Similar clustering was observed for segment M1, although the proportion of variance for the two first principal components was only 50.81 (**Fig. 6B**). On the other hand, for segments L3 and M3, T1L clustered tightly with T1L-T3pol, T3D clustered tightly with T3D-T1pol, and the proportions of variance for the two first principal components were 92.7 and 75.74, respectively (**Fig. 6C-D**). The PCA analyses quantitatively support more qualitative observations from the junction scatter plots and recombination frequency analyses and indicate that recombination profiles correspond with reovirus gene segment identity rather than that of the polymerase complex or other viral proteins.

**Figure 6.**
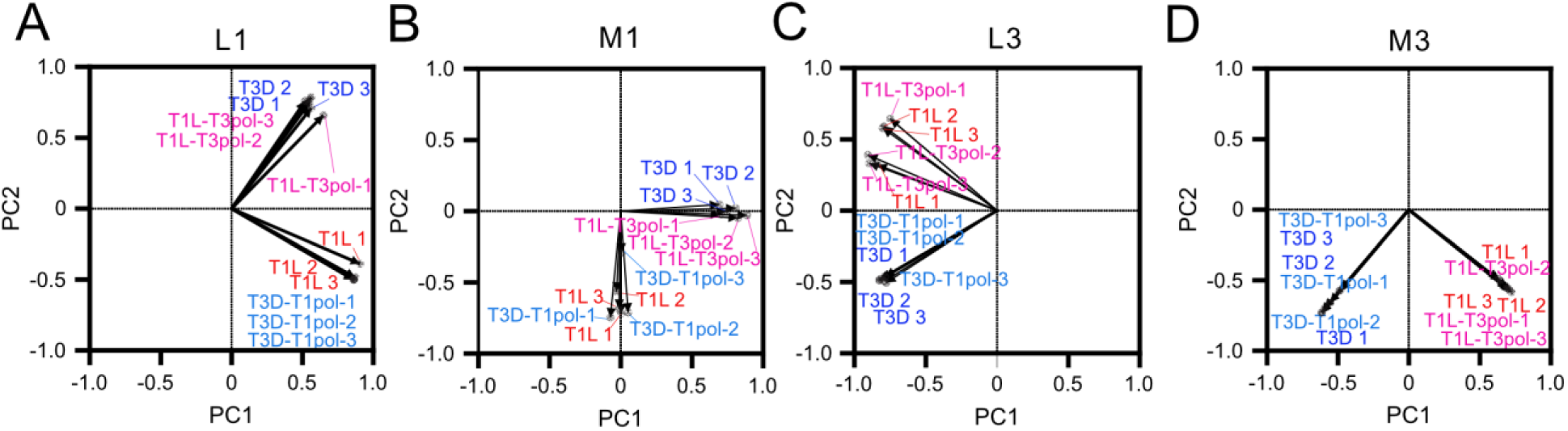
Principal component analysis of reovirus recombination profiles. Score plots for principal component analyses (PCA) of gene segments L1 (A), M1 (B), L3 (C), and M3 (D) applied to junctional recombination frequency data for three independent samples of reovirus strains T1L, T1L-T3pol, T3D-T1pol, and T3D.

### Recombination sites are conserved based on segment identity

Although PCA revealed that recombination junction profiles are similar for viruses that share a gene segment, a small number of overrepresented junctions may drive these similarities. Therefore, we determined the number of unique recombination junctions shared between reoviruses T1L, T1L-T3pol, T3D-T1pol, and T3D. We compared recombination junctions among all viruses for segments L1, L3, M1, and M3 (**Fig. 7**). For segments L1 and M1, T1L and T3D-T1pol or T3D and T1L-T3pol, in which these gene segments are identical, share many unique recombination junction sites. In contrast, T1L and T1L-T3pol or T3D and T3D-T1pol, in which these gene segments are of different origins, share few recombination junctions. For segments L3 and M3, the same is true of the reverse pairings: T1L and T1L-T3pol or T3D and T3D-T1pol share many junctions, while T3D and T1L-T3pol or T1L and T3D-T1pol share few junctions. Some common recombination junctions are anticipated among all strains for segments that share high sequence identity. However, like the results for recombination junction frequency and location, these observations suggest that recombination events resulting in DVG synthesis or packaging selection are predominantly determined by the identity of a reovirus gene segment rather than that of the polymerase complex or other viral proteins. Importantly, many recombination junctions, not just a small number of overrepresented junctions, are shared among reoviruses with similar recombination patterns.

**Figure 7.**
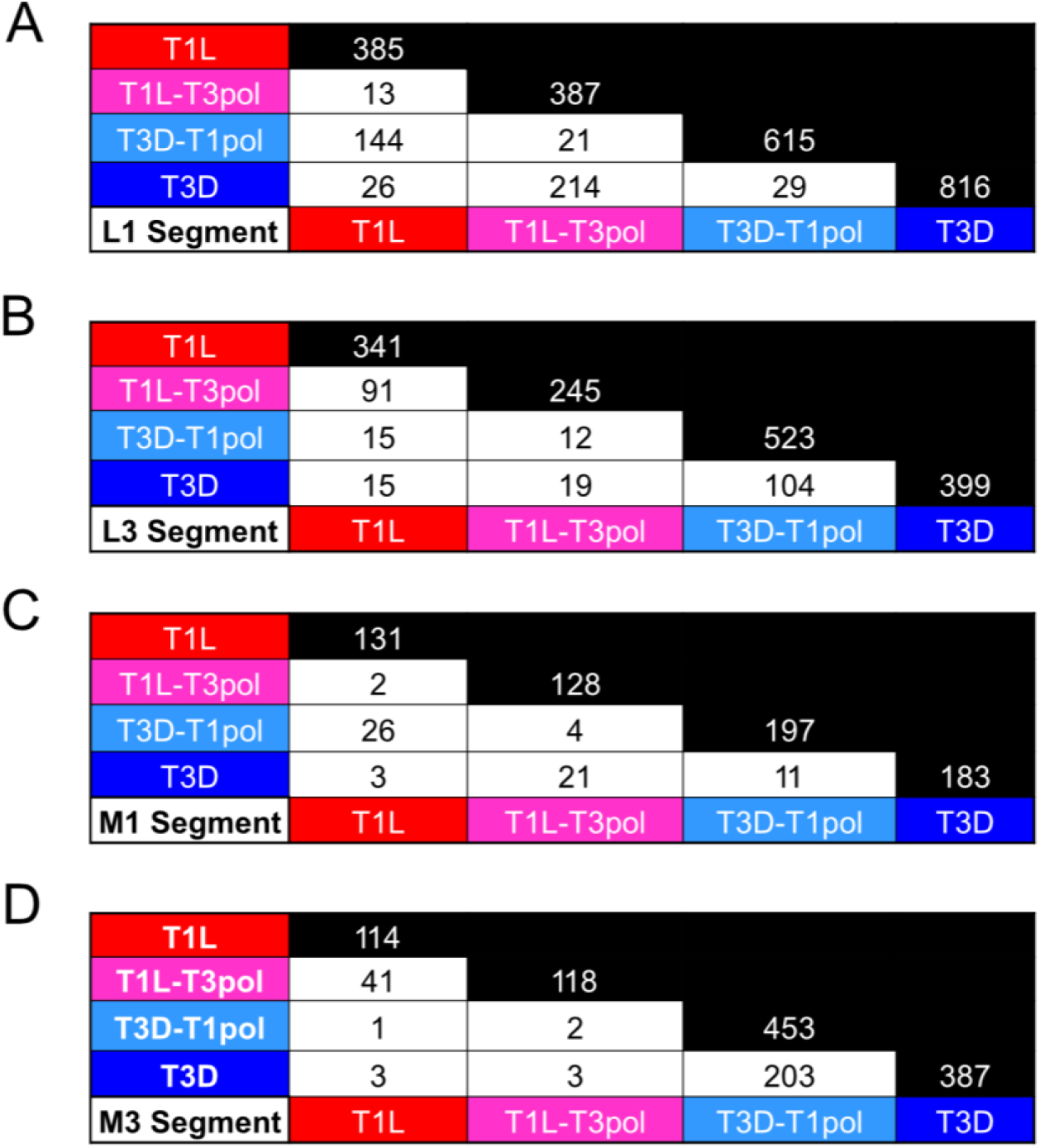
Recombination junctions shared among wild-type and polymerase-exchanged viruses. All the unique recombination junctions that are common between sequenced viral RNA from T1L, T1L-T3pol, T3D-T1pol, and T3D were identified using the “ggvenn” package in R. Analyses were conducted for segments L1 (A), L3 (B), M1 (C), and M3 (D). Black cells show total unique junctions for each virus within the indicated gene segment. Combined data from three independent samples of each virus were included in the analysis.

### Recombination events match nucleotides positionally between junction start and stop sites, but no specific motif is detected

For multiple virus families, the start and stop sites of recombination junctions have been associated with a specific nucleotide composition or with regions of microhomology (41, 48–51). For reovirus, regions of sequence microhomology, typically identical sequences about four to five nucleotides in length, appear to surround recombination junction sites (8). However, only two samples were investigated for the T1L strain using a sequencing method that can introduce artifactual recombination events. To validate the prior finding and determine whether regions of microhomology are also associated with recombination junction sites for T3D and polymerase-exchanged reoviruses, we analyzed all noncanonical junctions in segments L1 and L3. In each case, 50 nucleotides surrounding junction start sites and stop sites were analyzed. Every nucleotide surrounding the junction start site was matched to the equivalent position relative to the junction stop site. Positionally identical nucleotides were counted and graphed. For all analyzed segments and viruses, the nucleotides surrounding the junction start were frequently identical to the nucleotides surrounding the junction stop site (**Figs. 4A and 8A).** Most frequently, identical nucleotides were identified in about six positions surrounding recombination junction sites. For segment L1, substantially higher numbers of recombination junctions were detected in the negative strand than the positive strand, with high identity at the same relative positions (**Fig. S20A**). For segment L3, modestly more recombination junctions were detected for the positive strand than the negative strand (**Fig. S21A**). The percentages of junctions containing four, six, or eight identical nucleotides at junction sites were calculated for segments L1 and L3. For both segments, ∼ 10% of junctions for each virus contained eight identical nucleotides at the junction site, ∼ 40% contained six identical nucleotides, and ∼ 70-75% contained four identical nucleotides at the junction site (**Table. 2**). These findings suggest that recombination for all tested reoviruses, regardless of gene segment or polymerase complex identity, preferentially occurs at sites of microhomology. Thus, reovirus recombination is likely sequence-directed.

**Figure 8.**
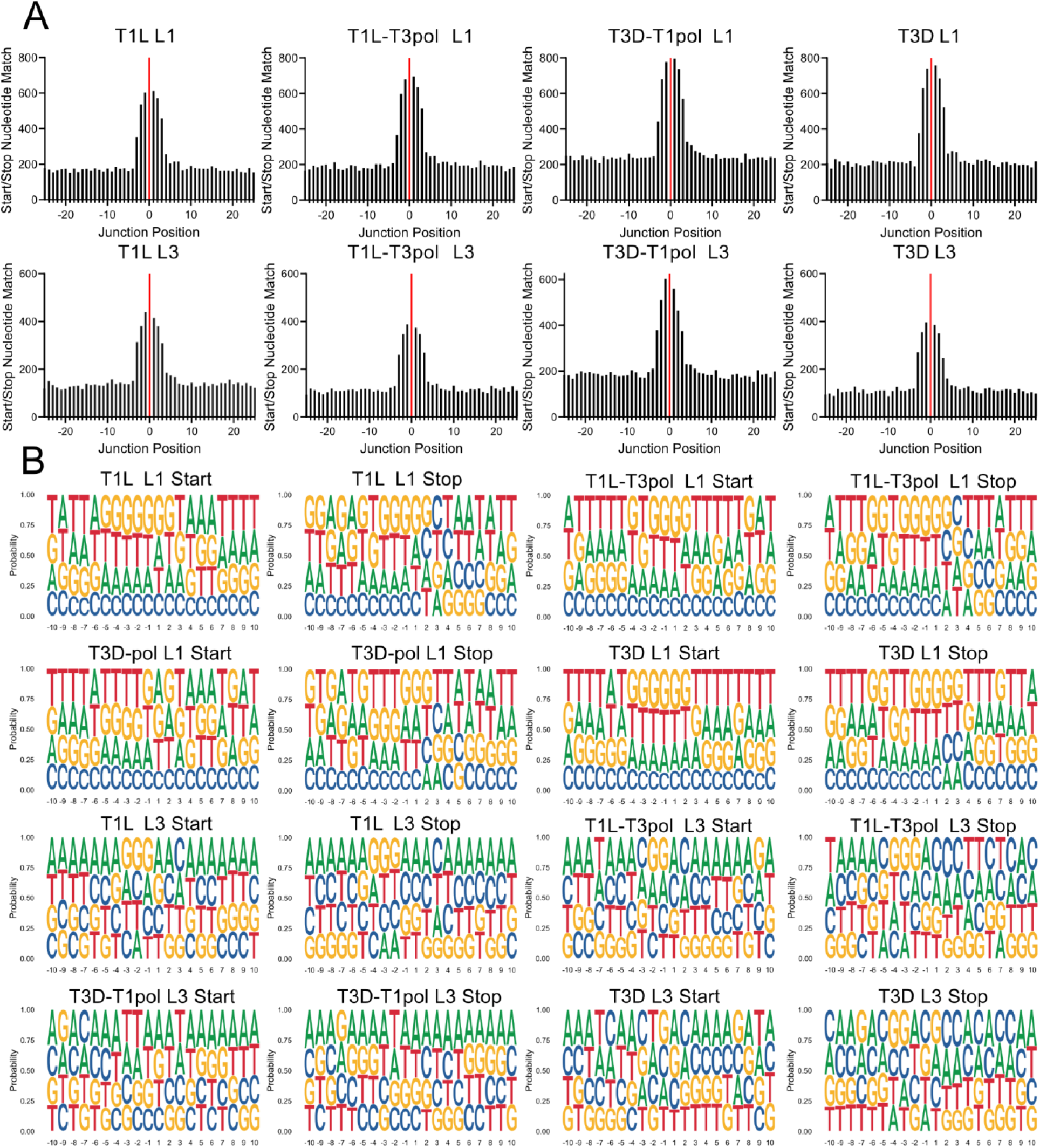
Microhomology and logo plots. (A) Sequence microhomology for T1L, T1L-T3pol, T3D-T1pol, and T3D. The nucleotides that overlap at each position in 25-bp regions flanking the start and stop recombination junction sites were quantified for segments L1 and L3. The junction is labeled zero and denotated with a solid red line. (B) Nucleotide composition was calculated as the percent adenosine (A), cytosine (C), guanine (G), and Thymine (T) at each position in a 10-bp region surrounding the start and stop recombination junction sites. Positions upstream (−10 to −1) and downstream (+1 to +10) of the junction position are indicated. Each analyzed virus contains combined data from three independent samples.

As nucleotides tend to match between recombination junction start and stop sites, it is possible that specific sequence motifs direct recombination. Therefore, nucleotide frequency relative to recombination junction start and stop sites was calculated for noncanonical junctions in segments L1 and L3. For all generated logoplots, no specific nucleotides were observed at a substantially higher frequency relative to the recombination start or stop site for any virus on either the positive or negative strand for either analyzed segment (**Figs. 8B and S20B-S21B**). Thus, there was no indication of a specific sequence motif or absolute nucleotide identity driving recombination. Instead, only positional identity relative to the junction start and stop site for a given recombination event is conserved.

### Recombination site selection is specific

After observing that recombination frequently occurs at start and stop sites containing a stretch of four identical nucleotides (**Fig. 8A and Table 2**), we wondered how frequently these stretches of identical sequence could be found in a given reovirus gene segment. We identified all the unique four-nucleotide sequences with microhomology at start and stop sites from recombination events in the L1 and L3 genes. Then, we identified all the occurrences of these sequences in L1 and L3. We found that in both segments, such four-nucleotide sequences typically appeared between 10 to 20 times (**Fig. 9A**). Some four-nucleotide sequences appeared much more frequently; ATGG appeared 41 times in the shared L1 genes of T1L and T3D-T1pol and 44 times in L1 of T3D and T1L-T3pol. Others had fewer occurrences, such as CCCC, which appeared six times in the L1 genes of T1L and T3D-T1pol. We next wondered how many occurrences of each of these four-nucleotide sequences were part of a recombination junction in our data set, either at the start or stop of the junction. Most occurrences of the four-nucleotide sequences (79-87%) were not part of any junctions in our data set (**Fig. 9B**). We found that for segments L1 and L3, ∼ 15% and ∼ 13% of occurrences of a four-nucleotide sequence were used in a junction once, indicating their incorporation into a single unique junction. Approximately 4% of occurrences of a four-nucleotide sequence for L1 and 2% of occurrences for L3 were selected for recombination events more than once, indicating their incorporation into more than one unique junction. The L1 sequences most frequently used for recombination were GCGG and GGGA in T1L, which were associated with six unique junction start sites. For the L1 stop position, the TGGG sequence was part of nine unique junctions. In segment L3, TAAT in T3D-T1pol was part of seven unique junction start sites, while TAAG was part of eleven unique junction stop sites. Overall, while a single four-nucleotide sequence can be frequently detected in a given reovirus gene segment, typically, only rare and specific occurrences of this sequence are utilized during recombination. These observations suggest that reovirus recombination is highly selective.

**Figure 9.**
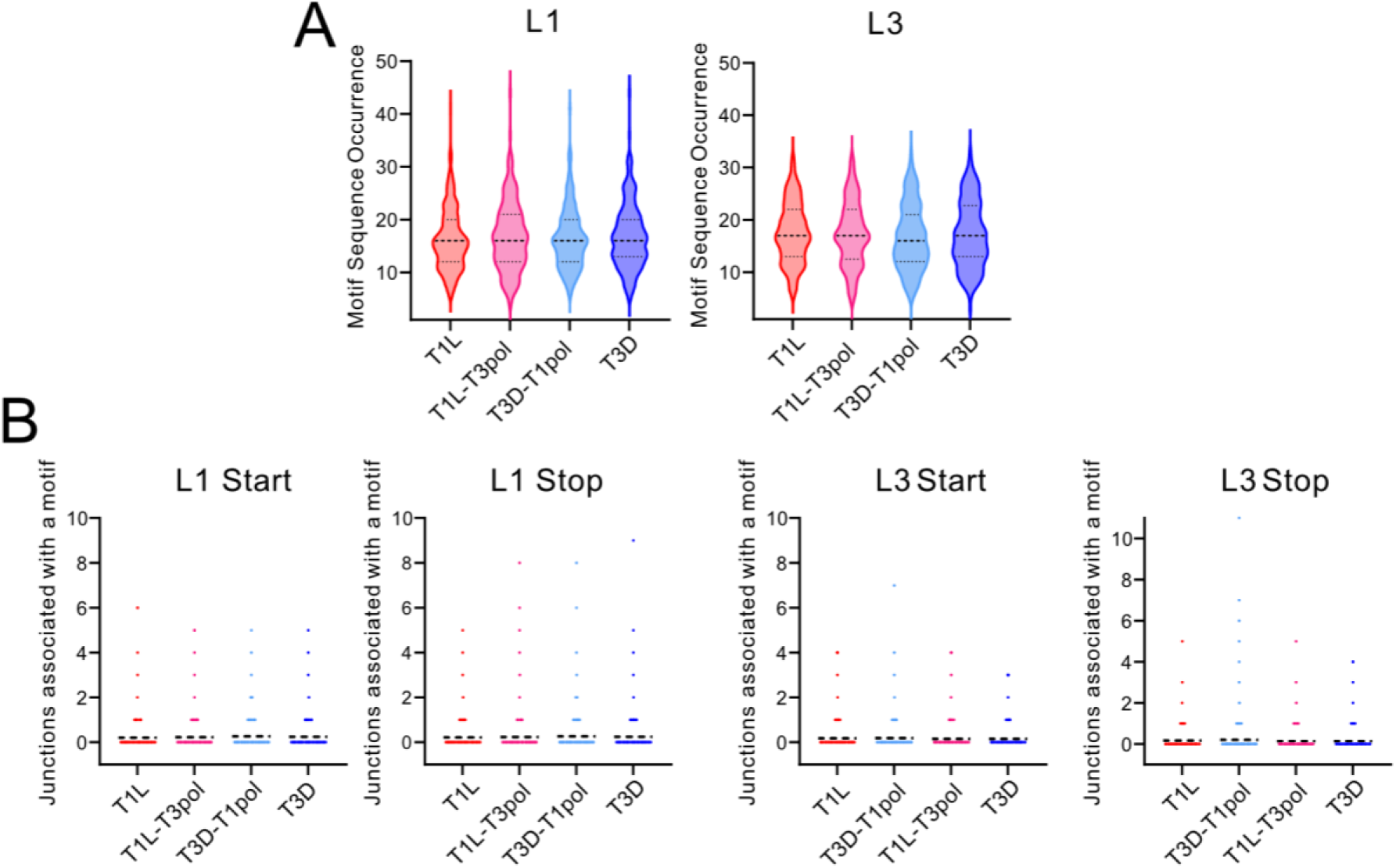
Recombination site selection. (A) Frequency of microhomology sequences in segments. Four-nucleotide sequences present in unique recombination junctions in the L1 or L3 segment from T1L, T1L-T3pol, T3D-T1pol, and T3D were identified. Then, the number of times each four-nucleotide sequence is present in segment L1 or L3 was computationally quantified. For L1, n = 176-240. For L3, n = 145-171. Dashed lines represent the median and quartiles. (B) Usage of microhomology sites for recombination. The number of times each occurrence of a four-nucleotide sequence from a unique start or stop site analyzed in (A) was incorporated into a recombination site in the L1 or L3 segment was computationally quantified for T1L, T1L-T3pol, T3D-T1pol, and T3D. For L1, n= 2915-3180. For L3, n= 2502-2851. Dashed lines represent the mean.

**Table 2.** Microhomology surrounding junctions within the L1 and L3 segments.

| L1 Segment |  |  |  | L3 Segment |  |  |  |
| --- | --- | --- | --- | --- | --- | --- | --- |
| Virus | Total Junctions | Number conserved | Percent Conserved | Virus | Total Junctions | Number Conserved | Percent Conserved |
| **Eight nucleotide positions conserved** |  |  |  |  |  |  |  |
| T1L | 663 | 65 | 10 | T1L | 525 | 41 | 8 |
| T1L-T3pol | 735 | 73 | 10 | T1L-T3pol | 430 | 48 | 11 |
| T3D-T1pol | 916 | 74 | 8 | T3D-T1pol | 709 | 63 | 9 |
| T3D | 800 | 54 | 7 | T3D | 420 | 57 | 14 |
| **Six nucleotide positions conserved** |  |  |  |  |  |  |  |
| T1L | 663 | 294 | 44 | T1L | 525 | 201 | 38 |
| T1L-T3pol | 735 | 297 | 40 | T1L-T3pol | 430 | 188 | 44 |
| T3D-T1pol | 916 | 330 | 36 | T3D-T1pol | 709 | 217 | 31 |
| T3D | 800 | 288 | 36 | T3D | 420 | 183 | 44 |
| **Four nucleotide positions conserved** |  |  |  |  |  |  |  |
| T1L | 663 | 506 | 76 | T1L | 525 | 333 | 63 |
| T1L-T3pol | 735 | 568 | 77 | T1L-T3pol | 430 | 316 | 73 |
| T3D-T1pol | 916 | 611 | 67 | T3D-T1pol | 709 | 394 | 56 |
| T3D | 800 | 600 | 75 | T3D | 420 | 323 | 77 |

## DISCUSSION

We sought to distinguish between contributions of RNA gene segments and the polymerase complex to virus strain-specific differences in DVG patterns. Although the presence of DVGs had been reported for reovirus (8, 44), the recombination mechanisms leading to DVG formation and their strain specificity remained unclear. We exchanged the polymerase complexes between reovirus strains T1L and T3D, which were previously shown to differ in their DVG patterns (**Fig. 1**) (8). RT-PCR profiling of DVGs for wild-type and polymerase-exchanged reoviruses failed to indicate a strict correlation between the polymerase complex and the diversity of DVG patterns among evolutionary lineages of serially passaged viruses (**Figs. 2 and S2-S4**). At least some DVGs carried in the virus population throughout the passage series shared identical sequences or internal deletions, suggesting a non-random recombination or packaging selection process (**Figs. 2**-**3, and S4**). Based on ClickSeq sequencing and computational analyses of packaged viral RNA, we found that T1L and T3D reoviruses have distinct recombination sites, which are conserved based on the identity of the gene segment rather than that of the polymerase complex (**Figs. 4-7 and S7-S19**). Regardless of segment or polymerase complex identity, we identified positional identity at recombination junction start and stop sites, where regions of microhomology were detected (**Figs. 8 and S20-S21**, **Table 2**). Although the conserved sequence at a recombination junction start and stop site were identified in many locations across a given gene segment, we found that recombination preferentially occurs only at a small subset of such locations (**Fig. 9**). Together these findings indicate that, like reovirus packaging, the recombination events that can yield DVGs may be highly orchestrated and primarily dictated by properties of viral RNA gene segments, which may include sequence and RNA secondary structure.

DVGs that arise during serial passage may affect the virus population and provide information about viral RNA synthesis and/or packaging requirements. In our analyzed reovirus series, some L1 and S4 DVGs persisted across all passages (**Figs. 2 and S4**). Other DVGs were transient, detected solely in a few passages, or prominent only in later passages. Variations in DVG diversity throughout passage series have been observed for other viruses such as SARS-CoV-2, herpesvirus, and Flock House virus (52–55). The persistence of a DVG in a virus population suggests that it is preferentially replicated and/or packaged (56). In most cases, there was little evidence of a substantial (>100-fold) decrease in viral titer across the series (**Fig. S1**), which suggests that most persistent reovirus DVGs detected are unlikely to be DIPs and may have neutral effects on the virus population. However, for segment L3, T3D-T1pol DVGs became more prominent as detection of the full-length segment decreased, which correlated with larger decreases in viral titers (**Figs. S1-S2**). DIPs have been identified for avian reovirus, for which S1 DVGs correlate with a reduction of full-length S1 and a decrease in infectious titers, suggesting a competitive relationship during replication and/or packaging (44). For the persistent DVGs of reovirus segments L1 and S4, the analyzed sequences revealed that these DVGs were identical between viruses or contained identical deletions (**Fig. 3**), which suggests that DVGs with specific characteristics are synthesized and/or packaged. Reovirus sequences located at the segment termini, which encompass the UTR and extend into the adjacent ORF, are thought to aid in assortment, packaging, and other replication processes (17–22, 57, 58). The DVGs that were sequenced retained the signals located at the 5′ end, but only a partial 3′ UTR. In a prior study, two of six sequenced DVGs from various segments also contained partial 3′ UTRs, although another two contained longer stretches of sequence at the 3′ end than the 5′ end (8). Further analysis of packaged DVGs could reveal the minimal components and features required for successful packaging.

In this study, a broad spectrum of recombination junctions was observed for T1L, T1L-T3pol, T3D-T1pol, and T3D reoviruses (**Figs. 4, S7-S10**). Based on RT-PCR profiling, DVG sequencing, and prior results, we expected to detect many small DVGs with large internal deletions (**Figs. 2, S2-S4**) (8, 44). Relatively few recombination events indicative of such DVGs were detected (**Figs. 4, S7-S10**). However, ClickSeq inefficiently captures products of ∼ 100-300 bp, limiting our capability to detect deletion DVGs that are missing considerable portions of a viral gene segment and to make correlations with results obtained from RT-PCR analyses of serially passaged viruses (36). Overall, our results identified junctions that indicate the existence of various packaged DVGs, including small and large internal deletions and duplications. Recombination events in the upper left corner of junction pattern plots, which represent the merger of a junction start site near the 3′ end of a segment with a stop site near the 5′ end of the same segment, are frequently discarded as artifacts. However, recombination events that generate duplications in viral gene segments have been identified for several dsRNA viruses [44-46, 60-64]. Based on ClickSeq data analyses alone, we cannot determine whether all RNA gene segments packaged by reovirus that contain recombination junctions also retain the segment termini, and we cannot completely define the packaged reovirus DVG catalog.

Several factors may influence the location of recombination sites in packaged reovirus RNA gene segments. In the analyses of ClickSeq data, distinct junctions and recombination sites were identified for T1L and T3D (**Figs. 4-5 and S7-S10**). These strains have high sequence identities in segments L1 (95.9%), L3 (97.7%), M1 (97.9%), and M3 (88.0%). Nonetheless, strain-specific recombination preferences were maintained in the presence of the polymerase complex of a different virus strain or upon encapsidation of the viral segment inside a different virus background. Recombination events leading to DVG generation have been observed to be enriched at specific regions of the viral genome for other RNA viruses, indicating preferred recombination sites (5, 59–61). Consistent with our results, rearrangements identified from rotavirus infections in independent immunocompetent children repeatedly occurred at a preferred location or hotspot (45–47). We also provide evidence that reovirus recombination occurs preferentially at specific sites of microhomology, short stretches of conserved sequence that are positionally identical between the recombination start and stop sites but it is not directed by a specific sequence motif (**Figs. 8 and S20-S21**). For rotaviruses, short regions of microhomology have been identified around the junctions of gene segment rearrangements (45–47). For SARS-CoV-2 and MERS-CoV, an enrichment of specific nucleotides upstream and downstream of the junction has been detected (41). Influenza A virus and Sindbis virus have been associated with specific sequence motifs in the deletion junctions that give rise to DVGs (48, 49). Thus, unrelated viruses also exhibit recombination specificity, although the overall mechanisms may differ. Additional factors that could influence recombination locations are RNA secondary structures or other RNA-RNA interactions. In SARS-CoV-2, it was found that secondary structures within the viral genome might guide recombination, as RNA structural distance between DVG junction positions was shorter than any two random genomic positions, which could facilitate the breaking and rejoining of those close pairs (5). It has also been shown that RNA secondary structure can influence chikungunya virus DVG production during replication (62). It has been predicted that the segment termini of +ssRNA segments of dsRNA viruses, such as reovirus and rotavirus, interact to form panhandle structures and contain stem-loops (58, 63–67). These predicted folds and structures are thought to be important for segment selection and packaging, but they may also be important for directing recombination. There is also evidence for intersegment RNA-RNA interactions between +ssRNAs during the assembly of dsRNA viruses including rotavirus and bluetongue virus (68–72). RNA secondary structure or other RNA-RNA interactions could influence the selection of the specific microhomology-containing regions that are used for recombination (**Fig. 9**). Determining the contributions of sequence and structure to DVG production will require the development of experimentally informed RNA secondary-structure and interaction models for +ssRNA reovirus segments to interrogate or model their influence on recombination events that might lead to DVG generation. Finally, packaging selection might influence our observations of recombination specificity. It is likely that a more diverse set of DVGs and recombination junctions is formed in infected cells than is selectively packaged into assembling viral particles.

Based on our new findings, we propose a model for the mechanism of RNA recombination in reoviruses (**Fig. 10**). The catalytic domain of reovirus RdRp λ3, like the RdRps of other dsRNA viruses, is sandwiched by N-and C-terminal domains that create a cage-like structure containing four tunnels for RNA template and product entry and exit (33, 34). This structure could not readily facilitate dissociation and reassociation of the polymerase on a distinct template. Therefore, a model of recombination that involves pausing and reinitiation at a different point along the same template is more likely. This model fits well with the detection of deletion DVGs, rearrangements, and recombination events anticipated to yield these types of noncanonical viral RNAs for dsRNA viruses (**Figs. 3, 4, and S7-S10**) (35, 44–47, 73, 74). The observation that ∼ 70% of the detected recombination junction sites occur at regions of microhomology, where positional identity was identified (**Table 2**), suggests that re-hybridization with the template might trigger the reinitiation of RNA synthesis. As dsRNA viruses undergo two RNA synthesis events, it is unknown whether the recombination events that lead to synthesis of reovirus DVGs and other noncanonical RNA molecules occur during transcription, in which +ssRNA is generated, or during genome replication, which produces the minus strand. For a segmented dsRNA ϕ6 bacteriophage, it has been shown that recombination can occur during minus-strand synthesis (75). It is highly unlikely that the dsRNA templates of transcription form substantial secondary structures or RNA-RNA interactions that might induce an RdRp pause. On the other hand, the +ssRNA templates of minus-strand synthesis (genome synthesis) are predicted to form complex secondary structures and RNA-RNA interactions that could influence RdRp activity or pausing, allowing the rehybridization of the template and reinitiation of RNA synthesis that could yield recombination sites in the viral genome (58, 63–67, 70, 76, 77).

**Figure 10.**
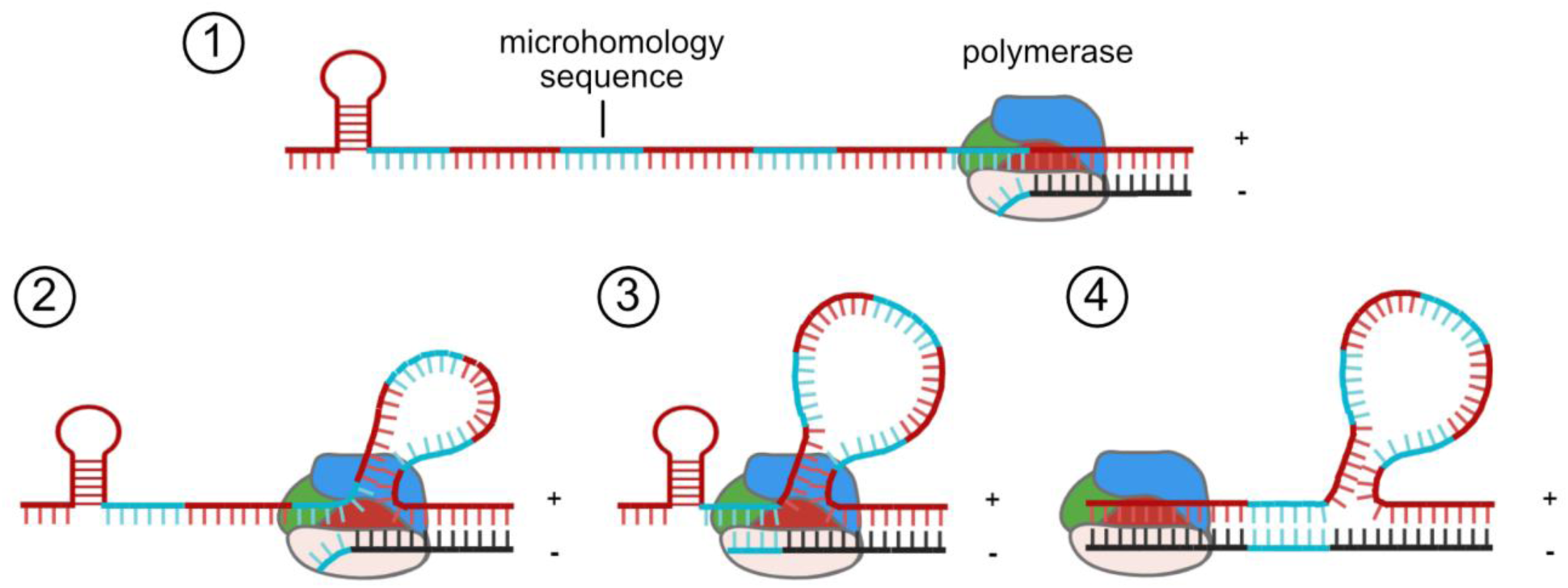
Recombination model for dsRNA viruses. (1) The RdRp pauses RNA synthesis at the junction start site during minus-strand synthesis. (2) The plus-strand template moves through the RdRp while synthesis is paused. Regions of sequence homology to junction start-site residues are bypassed. (3) The RdRp re-initiates synthesis at a specific junction stop site that has sequence homology to the junction start site. Annealing of the newly synthesized strand and other features, such as RNA secondary structure, interactions, or surrounding sequences, may influence re-initiation. (4) The RdRp continues synthesis to form a DVG or DVG template.

Observations in this study suggest that reovirus recombination occurs at preferred sites across the genome, which differ by reovirus strain. Rather than the polymerase complex, the identity of the gene segment seems to be a primary driver in selecting sites of recombination or the populations of DVGs that are packaged into reovirus particles. Together with properties of the reovirus RdRp and the types of reovirus DVGs detected, microhomology at recombination sites suggests a model of recombination in which the RdRp pauses synthesis and reinitiates after re-hybridization of the nascent product with the same template molecule at a different location (**Fig. 10**). Among the dsRNA viruses of the order Reovirales, many features are shared, including replication strategy, RdRp structure, packaging determinants, and noncanonical RNA properties. Thus, acquired knowledge for reovirus also may apply to the recombination mechanisms of other dsRNA viruses that affect human health (rotavirus), livestock (bluetongue), and agriculture (rice dwarf virus). Future studies should be aimed at determining associations of DVGs with pathogenesis, the influence of DVGs that are carried transiently or continuously in virus populations, discriminating between DVG synthesis and packaging in selecting the DVG population encapsidated by virus particles, defining the contributions of RNA sequence and structure to recombination, and determining precise recombination mechanisms for dsRNA viruses.

## MATERIALS AND METHODS

### Cells

L cells were maintained in suspension culture in Joklik’s minimum essential medium (JMEM; U.S. Biological) supplemented with 5% fetal bovine serum (FBS; Gibco), 2mM L-glutamine (Corning), and 100 U/ml penicillin plus 100 µg/ml streptomycin (Corning). Baby hamster kidney cells expressing T7 RNA polymerase controlled by a cytomegalovirus promoter (BHK-T7) cells were grown in Dulbecco’s minimum essential medium (DMEM; Corning) supplemented with 10% tryptose-phosphate broth, 1% nonessential amino acids, 5% FBS, 2 mM L-glutamine, and 100 U/mL penicillin plus 100 µg/mL streptomycin, with 1 mg/ml Geneticin (Gibco) added during alternate passages. MDCK cells were maintained in minimum essential medium with Earle’s salts (EMEM; Corning) supplemented with 10% FBS, 2 mM L-glutamine, and 100 U/ml penicillin plus 100 µg/ml streptomycin. All cells were cultivated at 37°C with 5% CO_2_.

### Viruses

Reovirus strains T1L, T3D, T1L-L1M1-T3D, and T3D-L1M1-T1L were engineered using plasmid-based reverse genetics (37, 38). T3D contained a T249I mutation that protects σ1, the attachment protein, from proteolytic cleavage (37). A pBacT7-S1T3D plasmid was previously engineered via ‘round the horn PCR to include the T249I mutation (78). T1L and T3D were rescued with a plasmid expressing C3P3 to increase efficiency (79). Briefly, monolayers of BHK-T7 cells at approximately 50% confluency in 12-well plates were co-transfected with 0.8 µg of pT7 plasmids encoding each of the ten T1L or T3D reovirus positive-sense viral RNAs and TransIT-LT1 transfection reagent (Mirus Bio LLC) (37, 38). For the polymerase complex-exchanged viruses, plasmids encoding T1L or T3D L1 and M1 positive-sense viral RNAs were introduced into the T3D or T1L reovirus background, respectively, in place of the native plasmids. Following five days of incubation, cells were subjected to two rounds of freezing and thawing. For viruses used in the passage series, lysates were amplified once in L cells, and viral titers were calculated by plaque assay in L cells (80). Viruses used for recombination analysis were plaque purified (80). Individual plaques were selected and amplified twice in L cells to generate stocks.

### Electropherotyping of recombinant reoviruses

4.8 × 10^6^ L cells were inoculated with three independently plaque-purified virus clones of each recombinant reovirus. Cells were incubated at 37°C for about five days until cytopathic effects were detected. Cells were collected and lysed in IGEPAL buffer (140 mM NaCl, 1.5 mM MgCl_2_, 100 mM Tris, pH 7.4) plus 5% IGEPAL CA-630 (Alfa Aesar). RNA from the supernatant was precipitated and resuspended in nuclease-free water. RNA samples were mixed with 4× sample buffer and heated for 10 min at 65°C. Samples were resolved by SDS-10% PAGE and ethidium bromide staining and visualized using a ChemiDoc MP imaging system (BIO-RAD).

### Replication kinetics analysis

To compare the replication kinetics of wild-type and polymerase-exchanged reoviruses, 8 × 10^5^ L or MDCK cells in 12-well plates were absorbed with three clones of each recombinant reovirus at an MOI of 0.1 PFU/cell. The unbound virus was removed by washing, and cells were incubated at 37°C for 0, 6, 12, 18, 24, or 48 h. After incubation, cells were lysed, and viral titer in lysates was quantified by plaque assay (80).

### Reovirus serial passages in L cells

To initiate the passage series, 5 × 10^7^ L cells were pelleted and adsorbed with T1L, T1L-T3pol, T3D-T1pol, T3D, or PBS (mock) in 10 ml JMEM at an MOI of 0.3 PFU/cell for 1 h at 37°C in triplicate from the same stock. Cell-virus mixtures were transferred into a suspension culture using 1 L bottles containing a magnetic stirring rod, and JMEM was added to a total volume of 100 ml. Suspension cultures were incubated for 48 h at 37°C, and the incubation was terminated by completing two freeze-and-thaw cycles and clearing debris from lysate by centrifugation at 1,000 × *g* for 10 min. For subsequent passages (P2 to P10), an identical protocol was followed, except that 5 × 10^7^ L cells were pelleted and resuspended in an adsorption inoculum of 10 ml of cleared lysate from the previous passage. For each virus, triplicate lineages were maintained throughout the series. Virus titer in each passage for each lineage was determined by plaque assay (80). One mock-infected passage was completed in triplicate.

### RT-PCR and electrophoretic analysis of DVG profiles

RNA was extracted from serial passage lysates using TRIzol LS Reagent (Invitrogen) according to manufacturer instructions. The RNA was used as a template with the OneStep RT-PCR kit (Qiagen) using manufacturer-recommended reaction conditions and gene-specific primers targeting the 5′ and 3′ termini to identify gene segments and DVGs. Primer sequences are available upon request. The extension time for segments >2 kb was 3 min and for those <2 kb was 1 min. RT-PCR products were resolved in 1.2% agarose stained with ethidium bromide and visualized using a ChemiDoc MP imaging system (BIO-RAD). RT-PCR products were resolved alongside a 100 bp DNA ladder and 1 kb DNA ladder (New England Biolabs).

### DVG sequencing

RT-PCR products from P6 and P9 present in multiple passages and lineages were gel extracted using a QIAquick gel extraction kit (Qiagen). The extracted DNA was submitted for Sanger sequencing (Azenta) with the primers used for RT-PCR amplification. Sequences were aligned to the corresponding reference viral genome using CLC sequence viewer 8 (Qiagen).

### Reovirus particle enrichment and RNA collection

4 × 10^8^ L cells were absorbed with three independent clones of stock for viruses T1L, T1L-T3pol, T3D-T1pol, and T3D at an MOI of 1 PFU/cell at 37°C for 72 h. Virus particles were purified from infected L cells by deoxycholate permeabilization, Vertrel XF (DuPont) extraction, and CsCl gradient centrifugation, as described previously(14). Reovirus particle concentration was determined from the equivalence of one unit of optical density at 260 nm to 2.1 × 10^12^ particles. 2.2 × 10^12^ viral particles for each individual virus and each replicate were treated with benzonase (250U) to degrade extra-particle nucleic acids, and the enzyme was inactivated with 0.5 M EDTA (pH 8.0). RNA was extracted from benzonase-treated particles using Trizol LS Reagent (Invitrogen), according to the manufacturer’s protocol. The quality of the RNA was determined using a Bioanalyzer (Agilent), and the RNA concentration was determined by Qubit (Invitrogen).

### ClickSeq library preparation and Next-Generation Sequencing (NGS)

ClickSeq libraries were generated from 250 ng of extracted RNA for each virus replicate. To initiate library preparation, the RNA underwent reverse transcription using SuperScript III reverse transcriptase (Invitrogen) with the addition of 3′-azido-nucleotides at a 1:35 AzNTPs:dNTPs ratio to induce chain termination and to release 3′-azido blocked cDNA fragments, which are click-ligated to Illumina i5 sequencing adaptors, as previously described (36). The products were PCR amplified with barcoded Illumina p7 adaptors, and libraries were sequenced by 150-bp paired-end sequencing on a NovaSeq X Plus sequencing system (Illumina). Assistance with quality control and next-generation sequencing was provided by ClickSeq Technologies LLC.

### Illumina RNA-sequencing processing and alignment

Raw reads were processed by first removing the Illumina TruSeq adapter using Cutadapt (81) with default settings. Reads shorter than 50 bp were discarded and reads containing any base with a PHRED score < 20 were removed using the FASTX toolkit. The raw FASTQ files were aligned to the T1L and T3D reovirus genome segments (NCBI or GenBank accession numbers AF378003.1, AF129820.1, AF461682.1, AF490617.1, AF174382.1, M14779.1, L19774.1, M14325.1, M13139.1, EF494435.1, EF494436.1, EF494437.1, EF494438.1, EF494439.1, EF494440.1, EF494441.1, EF494442.1, EF494443.1, EF494444.1; the T1L L1 sequence does not perfectly match those in GenBank and is available upon request) using the Python3 script *ViReMa* (Viral Recombination Mapper, version 0.32) (42) using the command line parameters python3 ViReMa.py reference_index input.fastq output.sam –Ouput_Dir sample_virema/--Output_Tag sample_virema -BED –MicroIndel_Length 5. The sequence alignment map (SAM) file was processed using the samtools (82) suite to calculate nucleotide depth at each position in a sorted binary alignment map (BAM) file (using command line parameters samtools depth -a -m 0 sample_virema.sorted.bam > sample_virema.coverage).

### Recombination junction analysis

Recombination junctions that contained micro insertions or deletions of 5 nt or less were filtered into separate output BED files. The frequency of the start and stop position of each junction was evaluated by dividing the frequency of reads mapping to a recombination junction by the total mapped reads. Junctions were plotted according to the genomic position and colored according to log_10_ of the frequency using *ggplot2* in RStudio (83). Recombination frequency was calculated at each genomic position by dividing the number of nucleotides in any junction mapping to the position by the total number of nucleotides sequenced at the position.

### Principal component analysis

A unique ID was created for each recombination event using the *dplyr* package in RStudio (84). All unique recombination events were identified for each virus, and the frequency value was added to a matrix. NA values were substituted with zero, and the data was normalized. Principal component analysis (PCA) was conducted and visualized for segments L1, L3, M1, and M3 using GraphPad Prism 10.3.1.

### Nucleotide composition analysis and sequence microhomology quantification

Nucleotide composition at each position surrounding recombination junction start and stop sites was determined. To avoid bias of highly replicated sites and to more closely reflect the stochastic nature of RNA recombination, each unique detected junction was counted equally rather than weighting by read count (54). Sequences were extracted from a sorted BED file that was converted into a text file to facilitate its manipulation. Start site and stop site sequences were separated in RStudio and read sequences containing less than 51 nucleotides in the stop site were filtered out. The nucleotide frequency at each position was calculated using the *Biostrings* package in RStudio (85). The nucleotide frequency was plotted as a logo plot using the *ggseqlogo* package in RStudio (86). For microhomology quantification, the nucleotides surrounding the junction sites were compared for positional identity relative to the start and stop sites. Positionally identical sequences were quantified in RStudio and plotted in GraphPad Prism 10.

### Microhomology sequence occurrence and usage

To calculate the number and percentage of recombination events containing microhomology regions of specific lengths, even numbers of nucleotides surrounding the junction sites (eight, six, or four) were extracted using *dplyr* (84). Extracted eight-, six-, or four-nucleotide sequences were compared between junction start and stop sites. Matches between the start and stop site were counted, and the percentage of junctions with microhomology were calculated using the *Biostrings* package in RStudio. To determine the frequency with which conserved four-nucleotide sequences found at recombination junctions recur in a given segment, unique four-nucleotide sequences with microhomology at start and stop sites from recombination events in the L1 and L3 genes were extracted using *dplyr* (84). For sequences originating from a negative strand, the reverse complement was obtained using *Biostrings* (*85*). Using the T1L or T3D L1 or L3 segment sequence as a reference, occurrences of each four-nucleotide sequence in the reference were quantified using *Biostrings* (*85*). The location of each short sequence in the viral genome was extracted using *Biostrings* (85). The location of short sequences from the negative strand and positive strand were adjusted to match the location of the start and stop sites of the recombination junctions detected by *ViReMa*. Then, the short sequence location number was counted for the number of occurrences in the column associated with the start and the stop sites separately from the dataset.

### Shared junction analysis

Individual recombination junctions were identified, and unique IDs were created for each recombination event using the *dplyr* (84) package in RStudio. Recombination junctions for all viruses within individual segments were compared to each other using the *ggvenn* package in RStudio (87), and numbers were plotted.

## ACKNOWLEDGEMENTS

We thank the Vanderbilt High Throughput Screening Facility for training and access to equipment. We thank Dr. James C. Slaughter for biostatistics expertise. Research reported in this work was supported by the National Institutes of Health (R01AI155646 to KMO). Alejandra Flores was supported by T32GM139800 and F31AI186483. The contents of this publication are solely the responsibility of the authors and do not necessarily represent the views of the National Institutes of Health.

## DATA AVAILABILITY STATEMENT

Data generated by ClickSeq sequencing can be accessed at NCBI Sequence Read Archive (SRA) under the BioProject accession PRJNA1259514. Code utilized in this report can be accessed at (https://github.com/ogdenlab1/ReoPolSwap.git).

