## Supplementary material for "Reovirus recombination is highly selective, and its profiles are primarily dictated by viral gene segment identity": Supplemetary data

### SUPPORTING INFORMATION

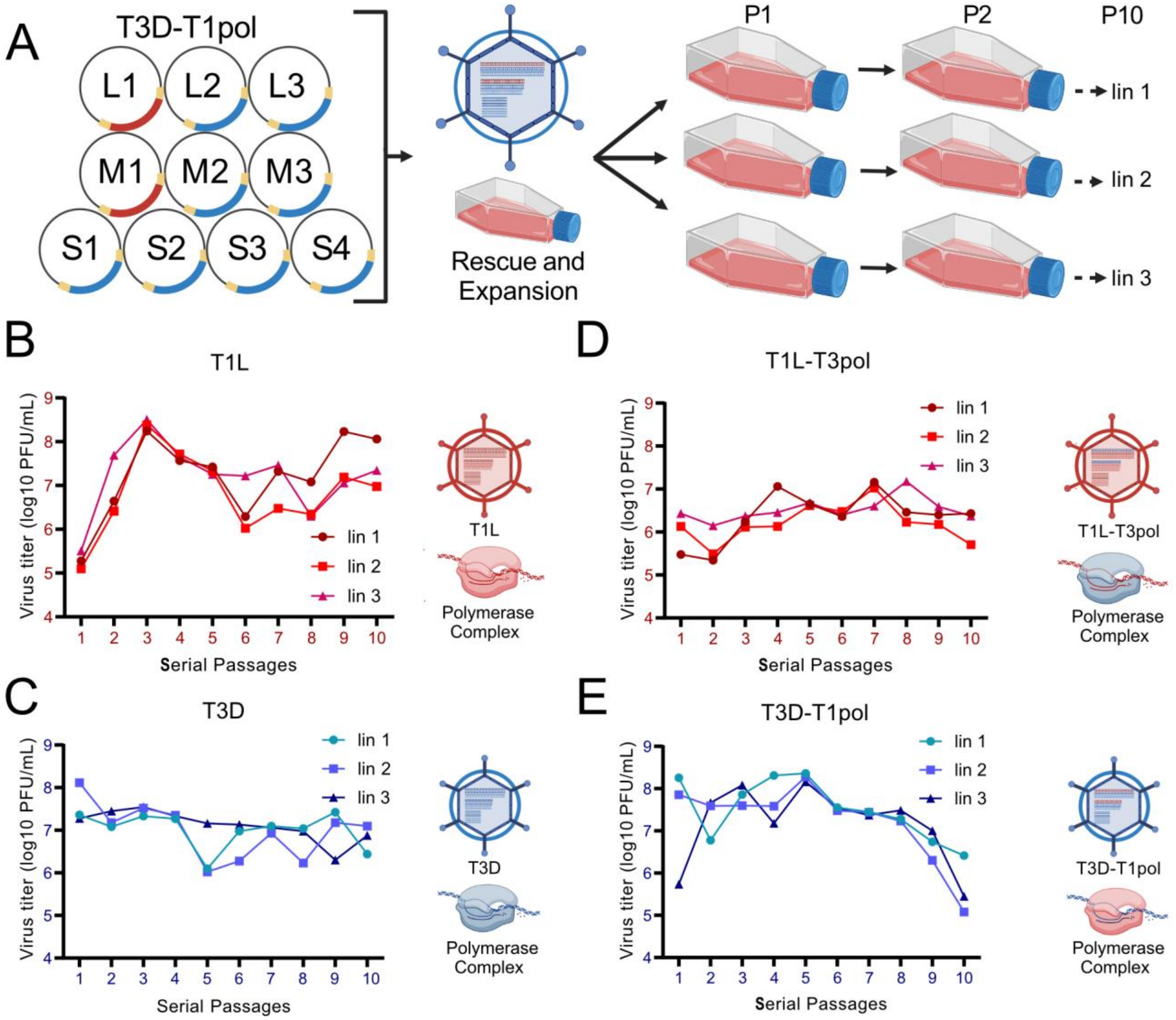


**Figure S1. Method and titers for serial passages of polymerase complex-exchanged reoviruses.** (A) On the left is a representation of the reverse genetics system used to generate wild-type reovirus strains or those containing exchanged L1 and M1 segments. Baby hamster kidney cells that express T7 RNA polymerase (BHK-T7) were transfected with 10 plasmids that encode positive-sense reovirus RNAs. Rescued reoviruses were amplified once in L cells and used as inocula for the first passage. L cells in suspension were adsorbed at an MOI of 0.3 PFU/cell in triplicate to compare evolutionary lineages and incubated for 48 h prior to lysis. Then, a new round of cells was infected with 10 ml of lysate from the previous passage. The procedure was repeated for 10 serial passages, as shown on the right. Created using Biorender.com. (B-E) Viral titers throughout serial passages were calculated by plaque assay for lysate from each of the three lineages of T1L (B), T3D (C), T1L-T3pol (D), or T3D-T1pol (E). Shown to the right of each graph is the origin of viral gene segments, represented as parallel lines inside the particle, the virus background, represented by the particle, and the polymerase complex for the passaged virus, indicated by color (red, T1L; blue, T3D). Particle and polymerase complex schematics created using Biorender.com.

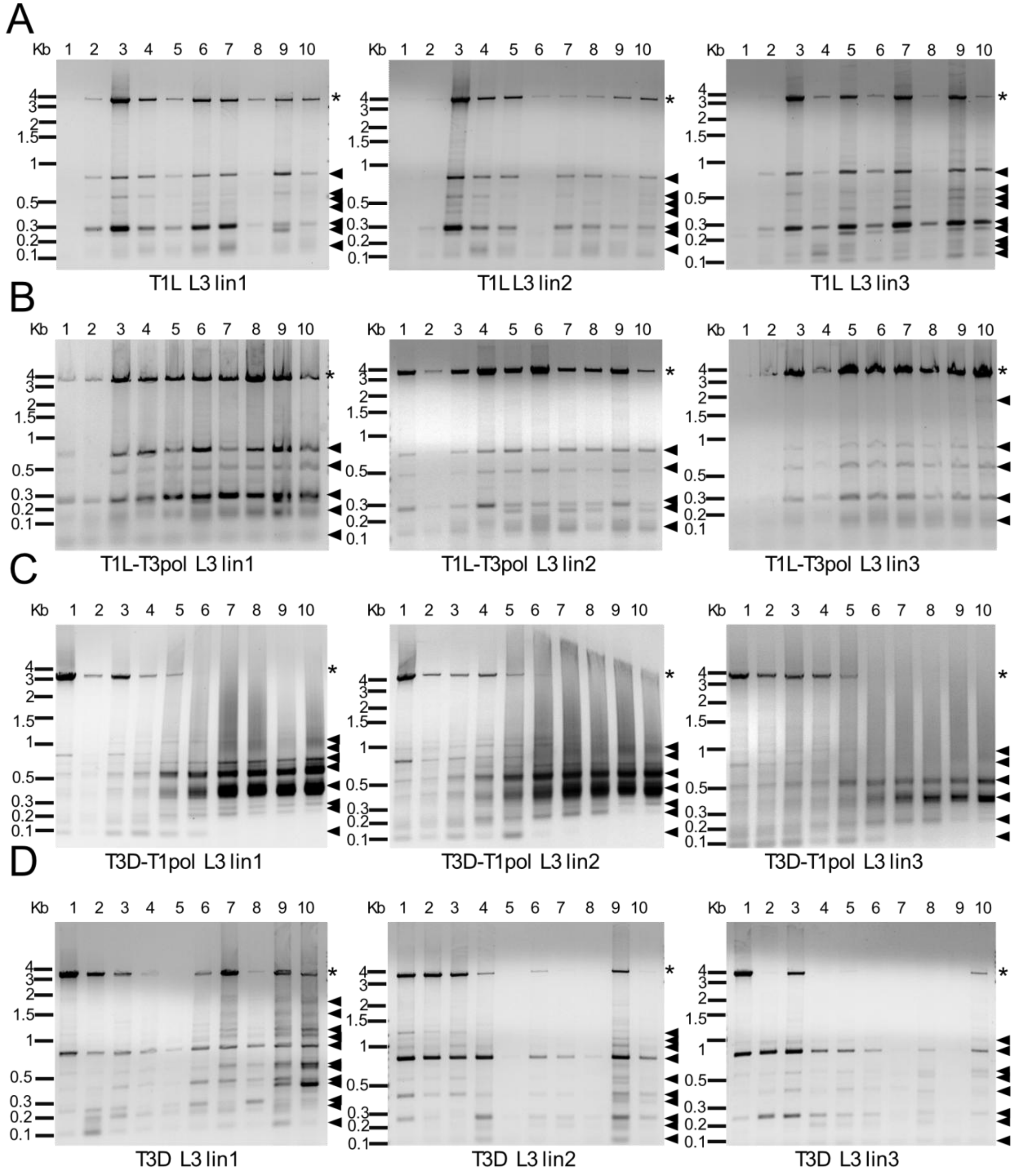


**Figure S2. Serial passage lineage 1 to 3 reovirus gene segment L3 patterns.** RNA extracted from serial passages was used as a template for RT-PCR reactions in which primers targeting terminal sequences in the L3 segments of both T1L and T3D were included. Products from T1L (A), T1L-T3pol (B), T3D-T1pol (C), and T3D (D) P1-P10 were resolved in 1.2% ethidium bromide-stained agarose gels. The evolutionary lineage is indicated. Black asterisks indicate the position of the full-length segment. Triangles indicate products smaller than the full-length segment.

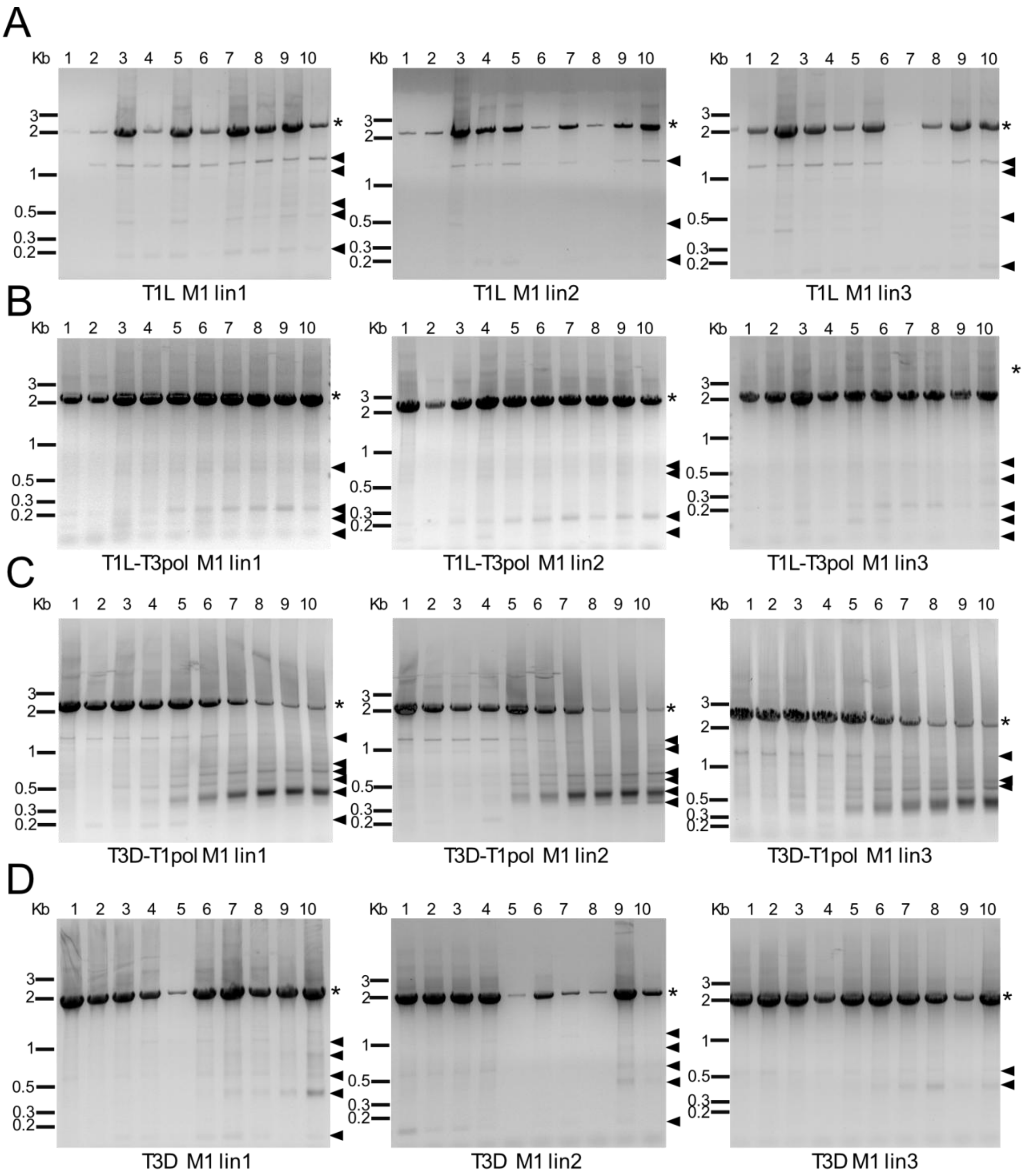


**Figure S3. Serial passage lineage 1 to 3 reovirus gene segment M1 patterns.** RNA extracted from serial passages was used as a template for RT-PCR reactions in which primers targeting terminal sequences in the M1 segments of both T1L and T3D were included. Products from T1L (A), T1L-T3pol (B), T3D-T1pol (C), and T3D (D) P1-P10 were resolved in 1.2% ethidium bromide-stained agarose gels. The evolutionary lineage is indicated. Black asterisks indicate the position of the full-length segment. Triangles indicate products smaller than the full-length segment.

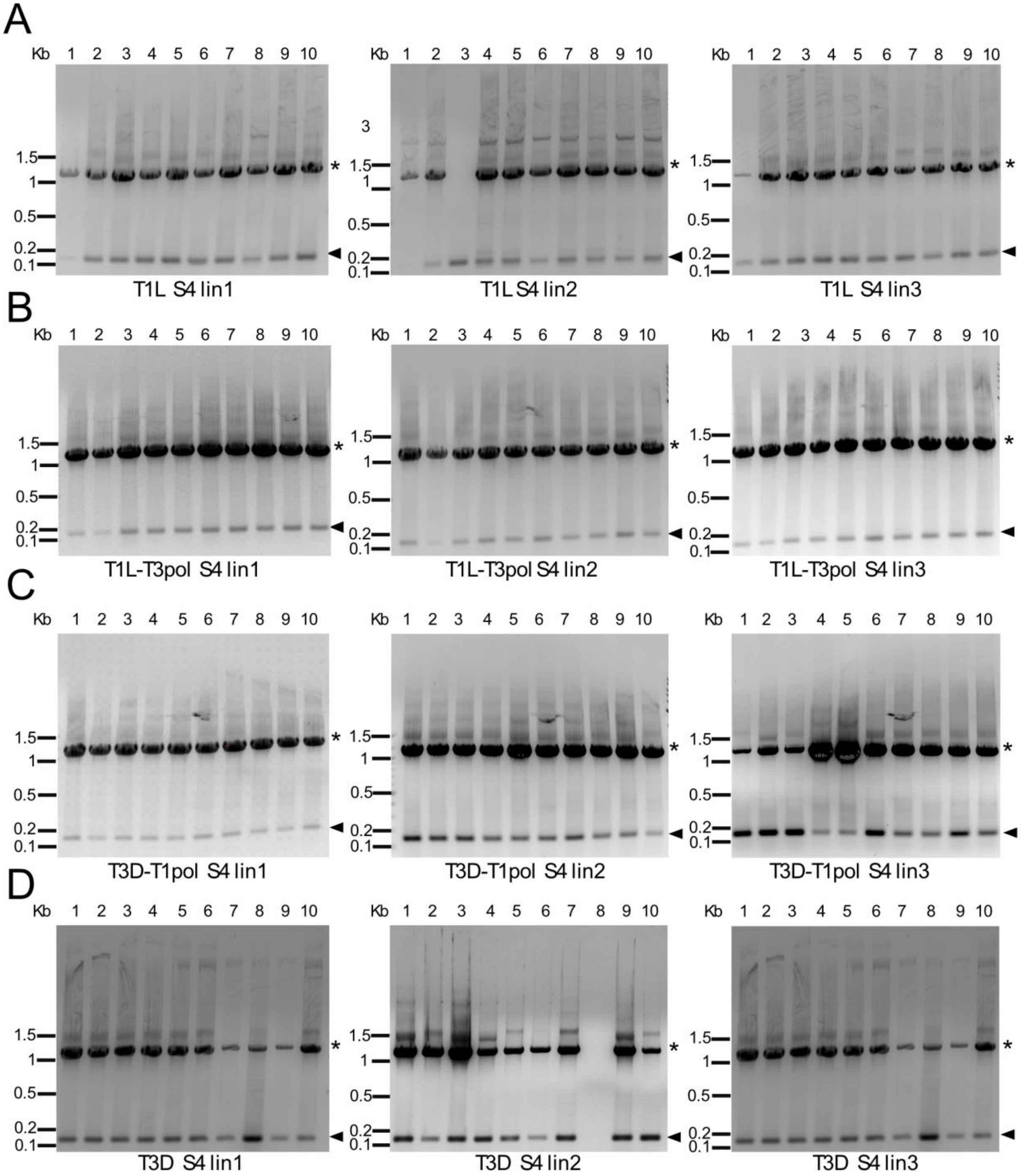


**Figure S4. Serial passage lineage 1 to 3 reovirus gene segment S4 patterns.** RNA extracted from serial passages was used as a template for RT-PCR reactions in which primers targeting terminal sequences in the S4 segments of both T1L and T3D were included. Products from T1L (A), T1L-T3pol (B), T3D-T1pol (C), and T3D (D) P1-P10 were resolved in 1.2% ethidium bromide-stained agarose gels. The evolutionary lineage is indicated. Black asterisks indicate the position of the full-length segment. Triangles indicate products smaller than the full-length segment.

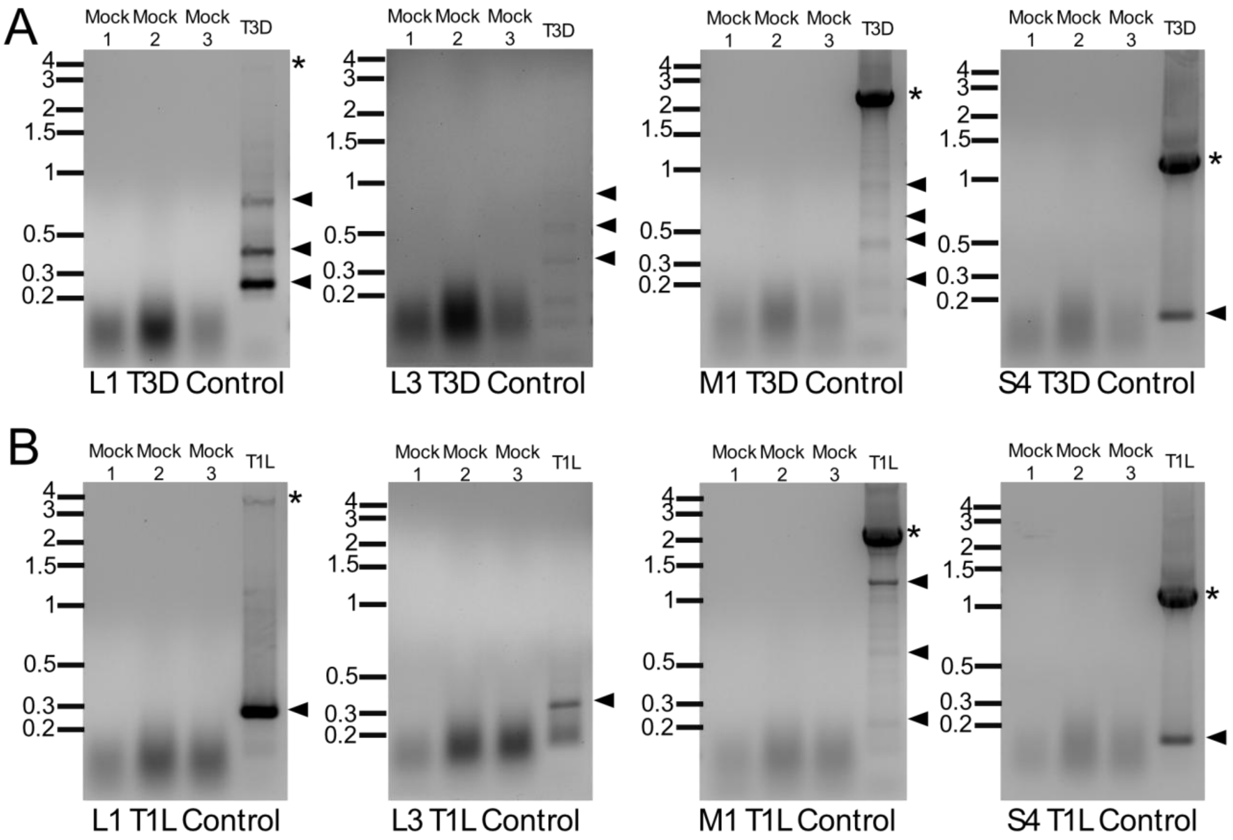


**Figure S5. Mock serial passage control RT-PCR patterns.** (A-B) RNA extracted from mock-infected L cells in triplicate and from T1L or T3D P9 lin1 was used as a template for RT-PCR reactions in which primers targeting terminal sequences in the L1, L3, M3, and S4 segments of both T1L and T3D were included. Products from mock controls and T3D (A) or from mock controls and T1L (B) were resolved in 1.2% ethidium bromide-stained agarose gels. The evolutionary lineage is indicated. Black asterisks indicate the position of the full-length segment. Triangles indicate products smaller than the full-length segment.

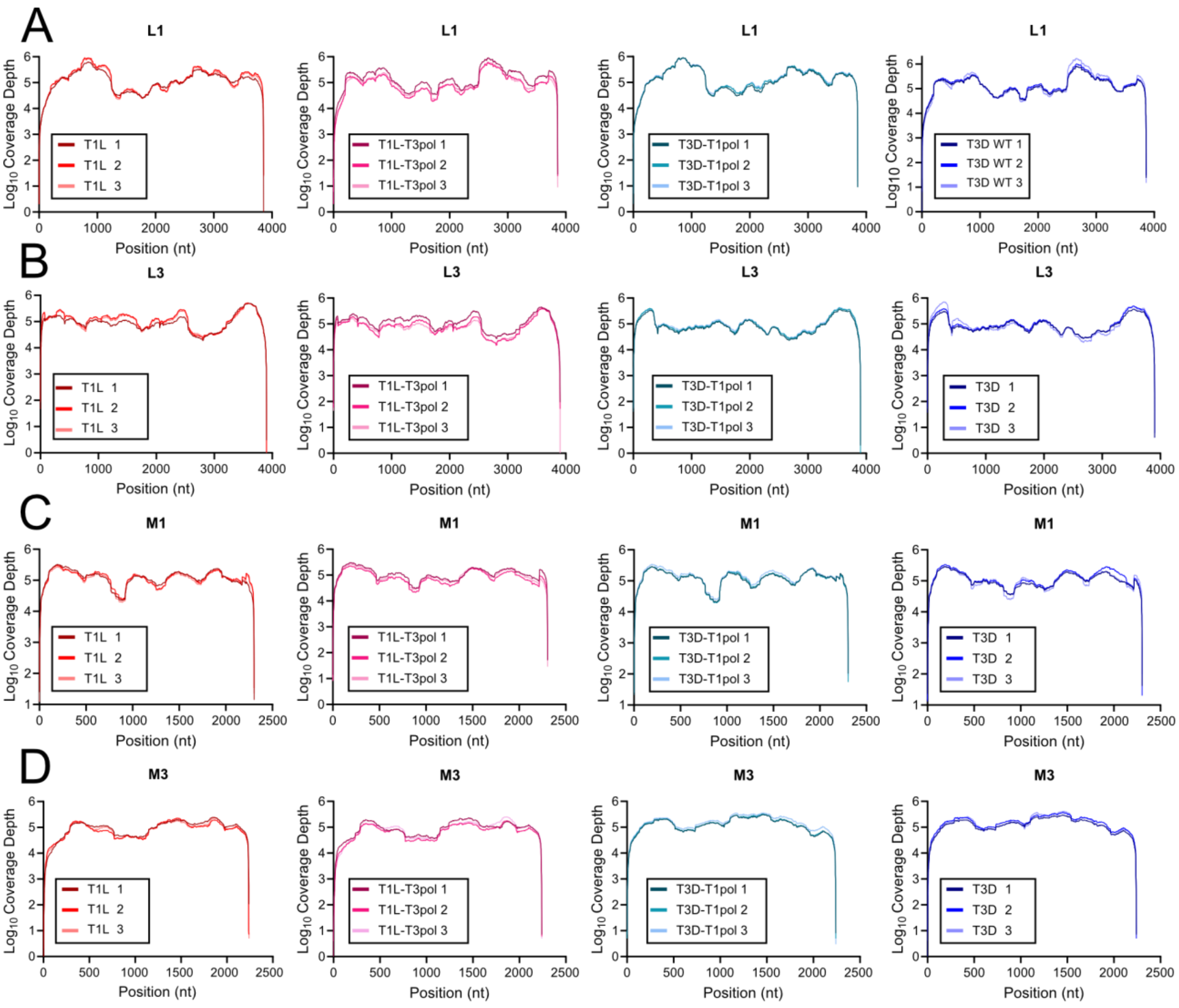


**Figure S6. Reovirus sequence coverage across genome segments.** Sequence read depth across all nucleotide positions for segments L1 (A), L3 (B), M1 (C), and M3 (D) of T1L, T1L-T3pol, T3D-T1pol, and T3D. Coverage for three individual virus clones is shown.

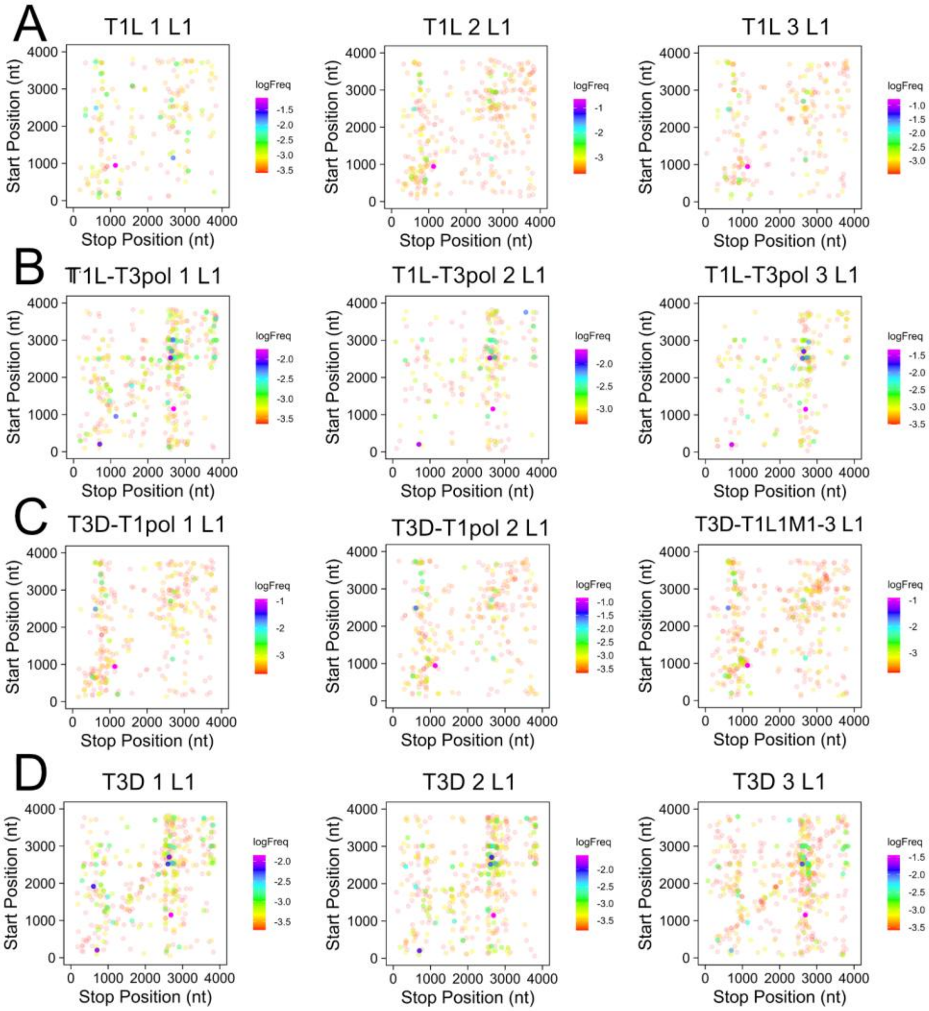


**Figure S7. Reovirus junction site location and frequency: Segment L1.** Recombination junction site location and frequency in sequenced T1L (A), T1L-T3pol (B), T3D-T1pol (C), and T3D (D) virion RNA for gene segment L1 in three independent samples. Junction sites are indicated by dots whose position corresponds to upstream and downstream sequences that are merged to form a novel junction. Junction frequency is indicated by dot color, according to the legend to the right of each image.

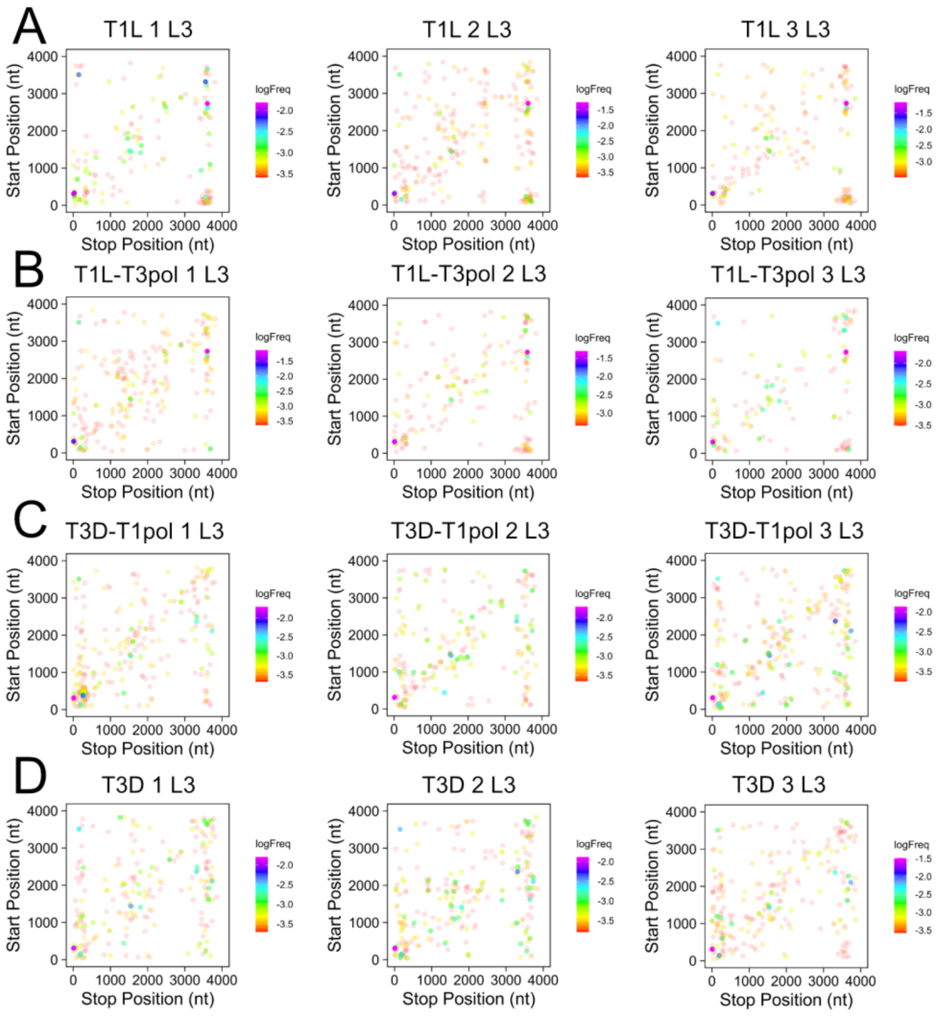


**Figure S8. Reovirus junction site location and frequency: Segment L3. Recombination** junction site location and frequency in sequenced T1L (A), T1L-T3pol (B), T3D-T1pol (C), and T3D (D) virion RNA for gene segment L3 in three independent samples. Junction sites are indicated by dots whose position corresponds to upstream and downstream sequences that are merged to form a novel junction. Junction frequency is indicated by dot color, according to the legend to the right of each image.

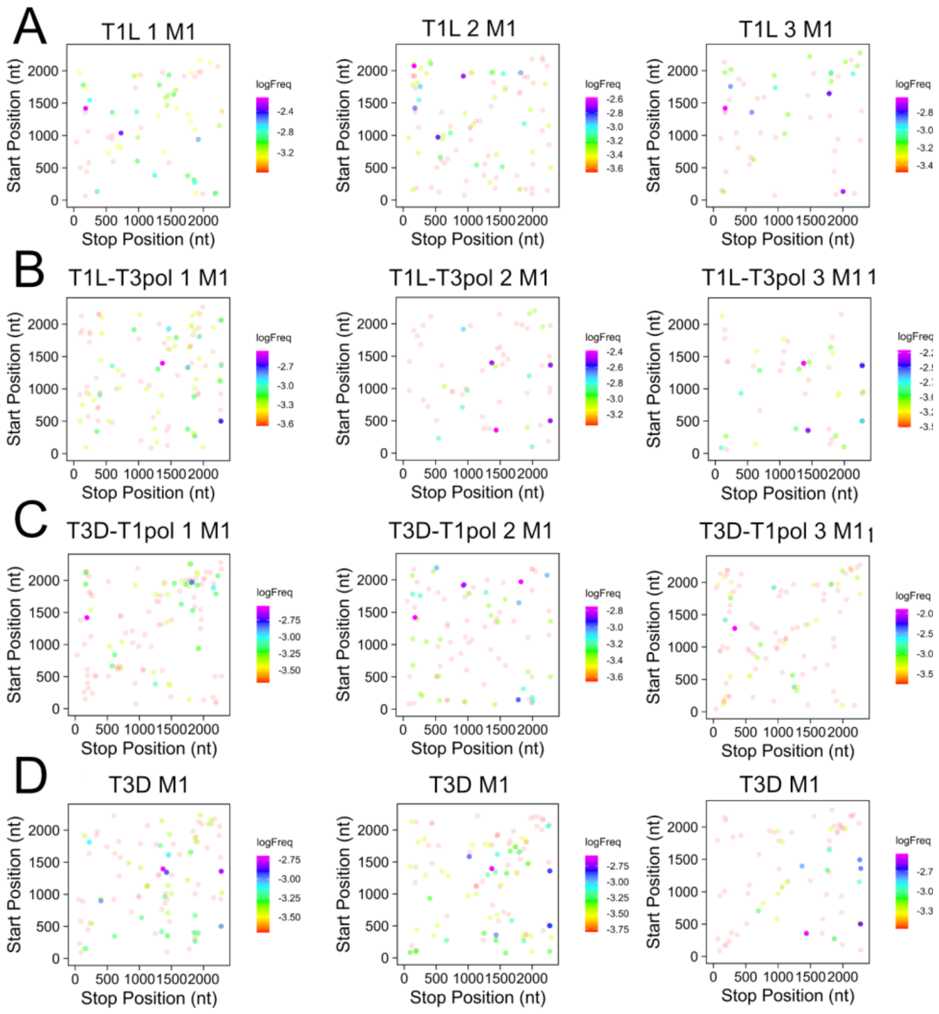


**Figure S9. Reovirus junction site location and frequency: Segment M1.** Recombination junction site location and frequency in sequenced T1L (A), T1L-T3pol (B), T3D-T1pol (C), and T3D (D) virion RNA for gene segment M1 in three independent samples. Junction sites are indicated by dots whose position corresponds to upstream and downstream sequences that are merged to form a novel junction. Junction frequency is indicated by dot color, according to the legend to the right of each image.

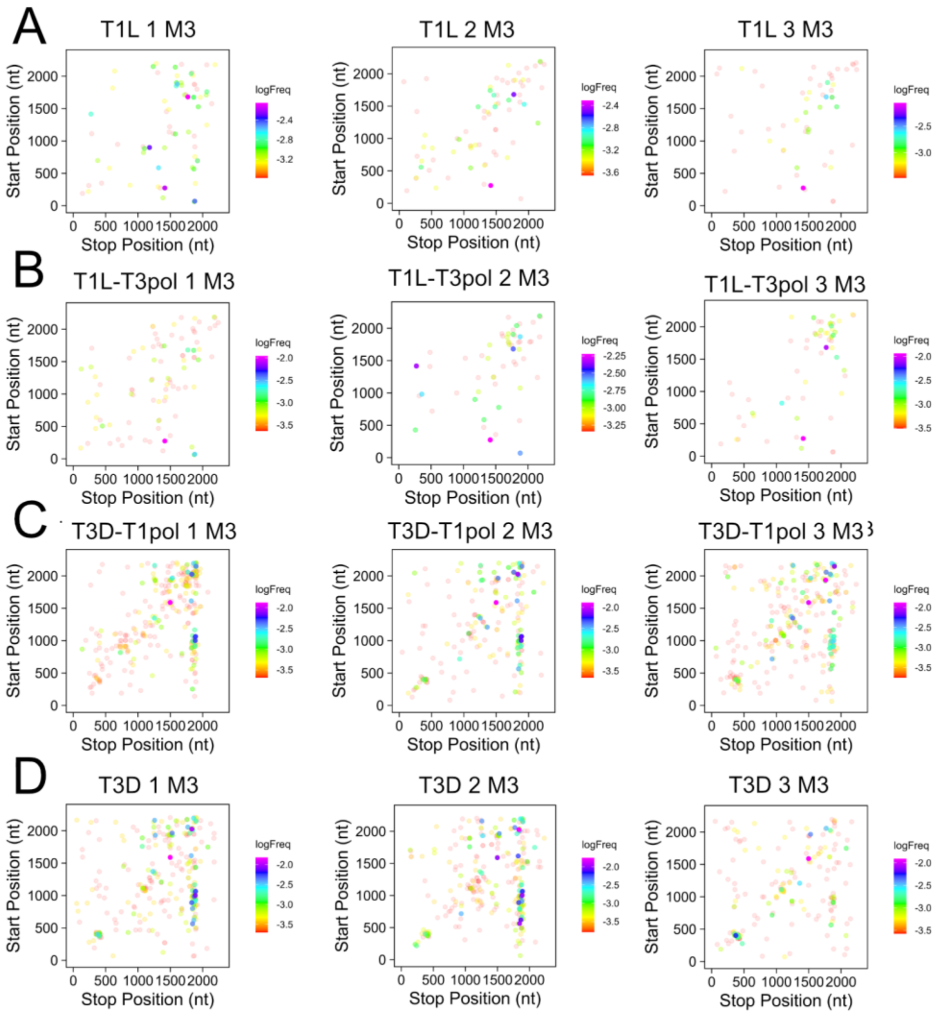


**Figure S10. Reovirus junction site location and frequency: Segment M3**. Recombination junction site location and frequency in sequenced T1L (A), T1L-T3pol (B), T3D-T1pol (C), and T3D (D) virion RNA for gene segment M3 in three independent samples. Junction sites are indicated by dots whose position corresponds to upstream and downstream sequences that are merged to form a novel junction. Junction frequency is indicated by dot color, according to the legend to the right of each image.

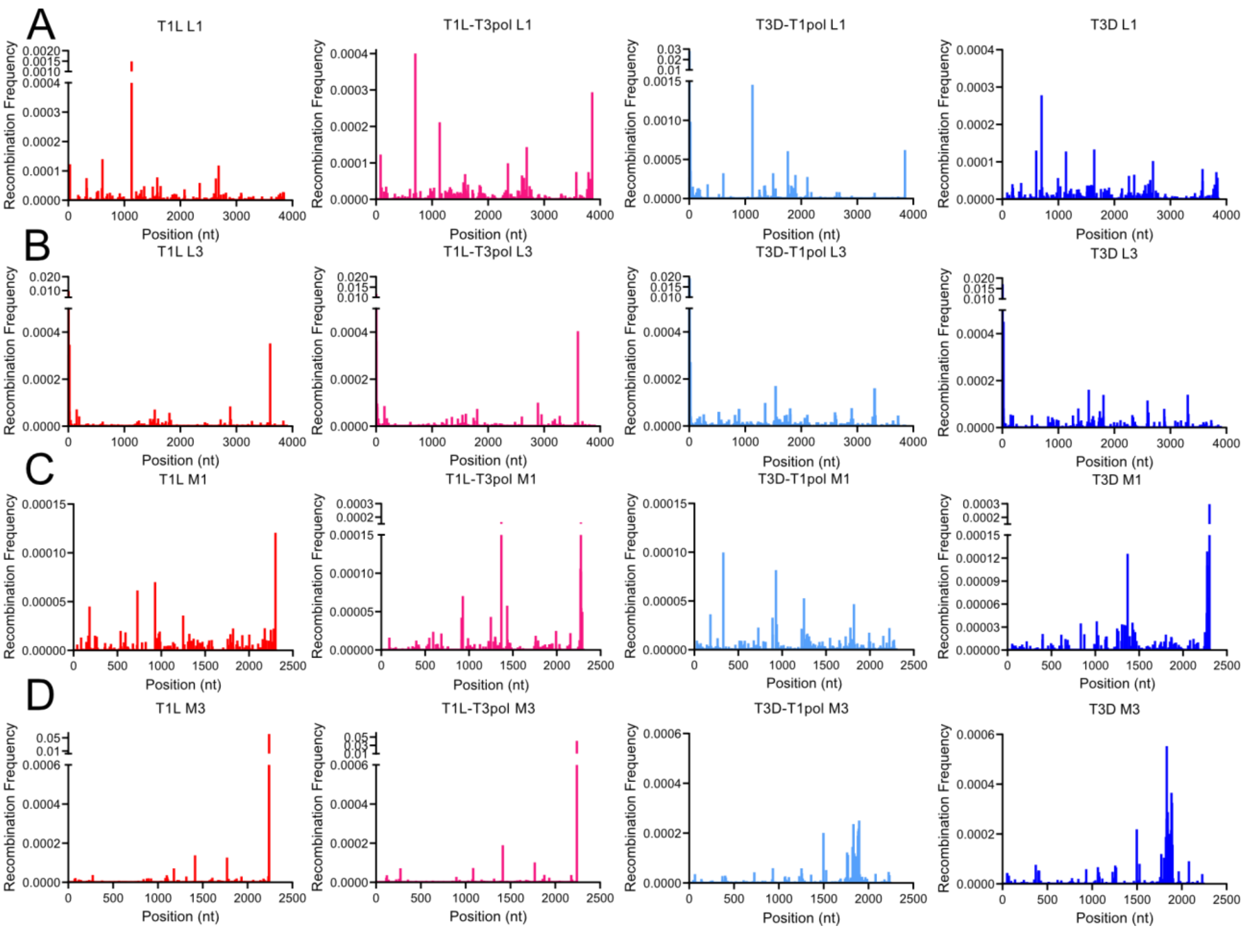


**Figure S11. Reovirus recombination stop-site frequency across gene segments**. Recombination stop-site frequency at each nucleotide position across L1 (A), L3 (B), M1 (C), and M3 (D) segments from T1L, T1L-T3pol, T3D-T1pol, and T3D. Mean positional recombination frequency for sequenced virion RNA from three clones per virus is shown.

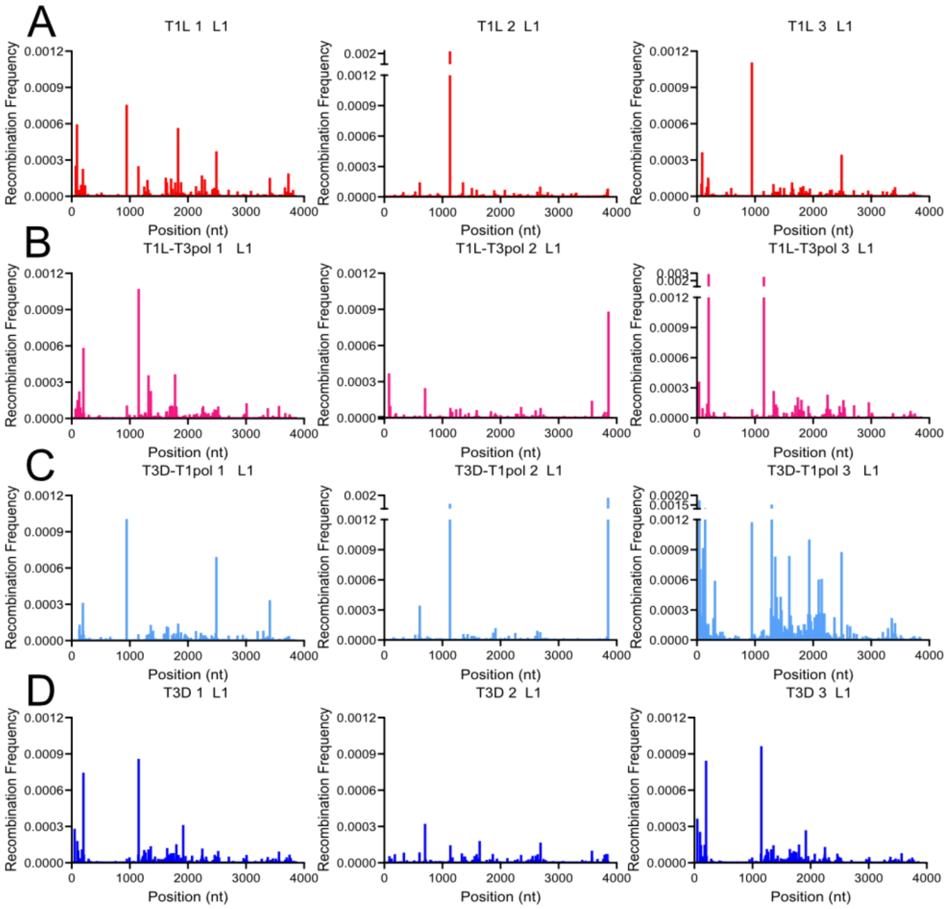


**Figure S12. Reovirus recombination start-site frequency across gene segments: Segment L1.** Recombination start-site frequency at each nucleotide position across the L1 segment from T1L (A), T1L-T3pol (B), T3D-T1pol (C), and T3D (D). Positional recombination frequency of sequenced virion RNA for each individual virus clone is shown.

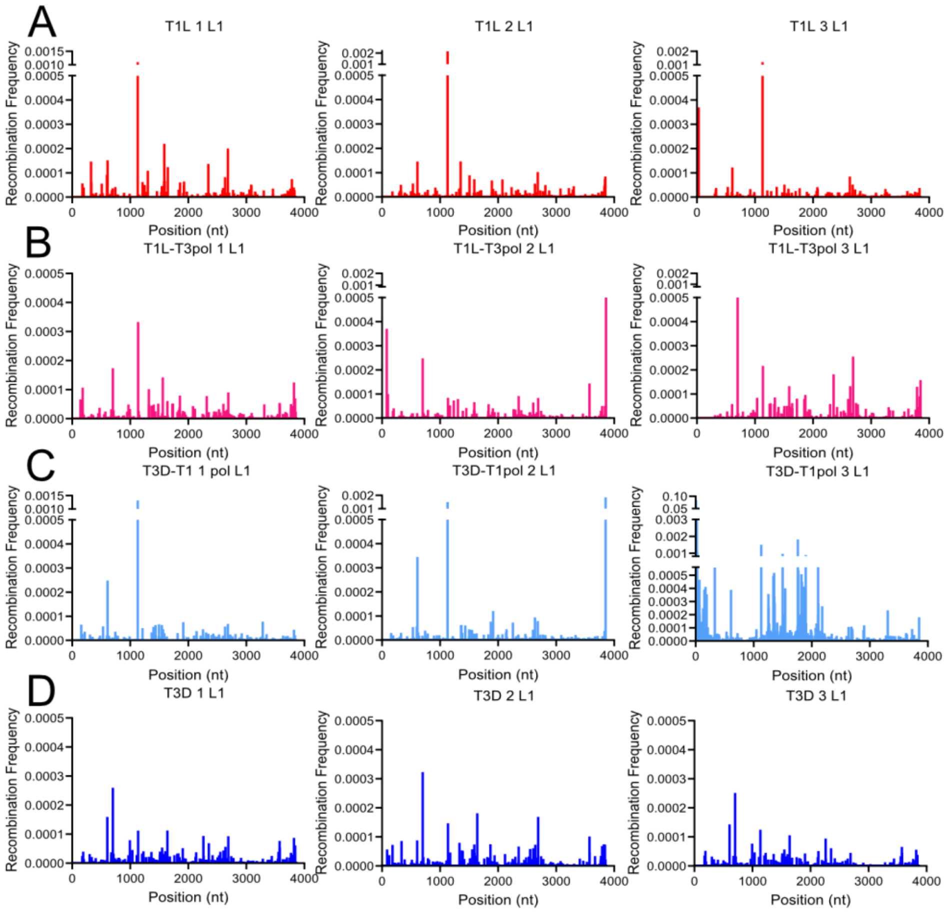

**Figure S13. Reovirus recombination stop-site frequency across gene segments: Segment L1.** Recombination stop-site frequency at each nucleotide position across the L1 segment from T1L (A), T1L-T3pol (B), T3D-T1pol (C), and T3D (D). Positional recombination frequency of sequenced virion RNA for each individual virus clone is shown.

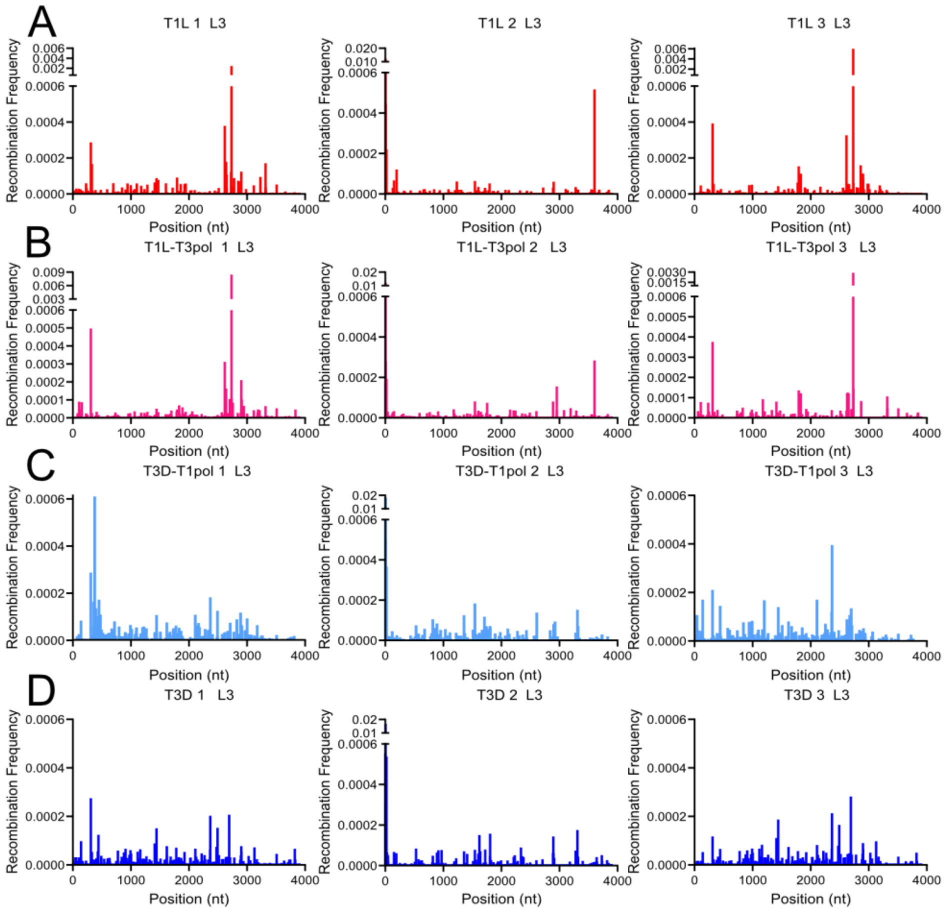


**Figure S14. Reovirus recombination start-site frequency across gene segments: Segment L3.** Recombination start-site frequency at each nucleotide position across the L3 segment from T1L (A), T1L-T3pol (B), T3D-T1pol (C), and T3D (D). Positional recombination frequency of sequenced virion RNA for each individual virus clone is shown.

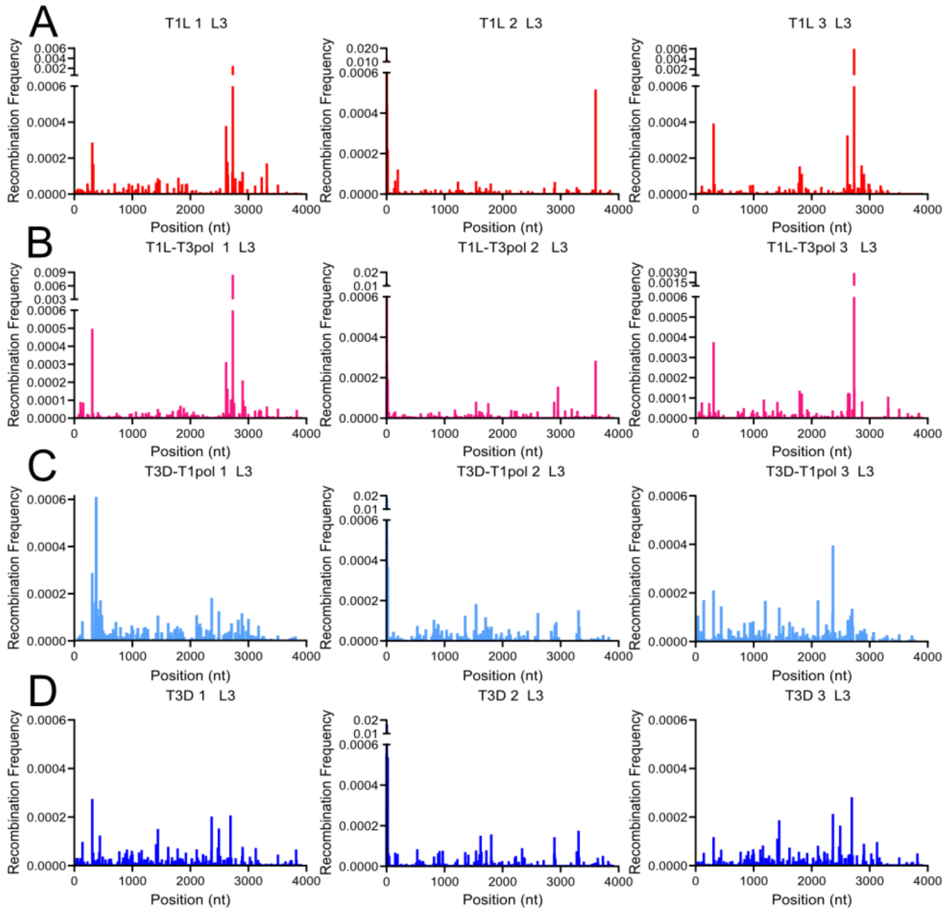

**Figure S15. Reovirus recombination stop-site frequency across gene segments: Segment L3.** Recombination start-site frequency at each nucleotide position across the L3 segment from T1L (A), T1L-T3pol (B), T3D-T1pol (C), and T3D (D). Positional recombination frequency of sequenced virion RNA for each individual virus clone is shown.

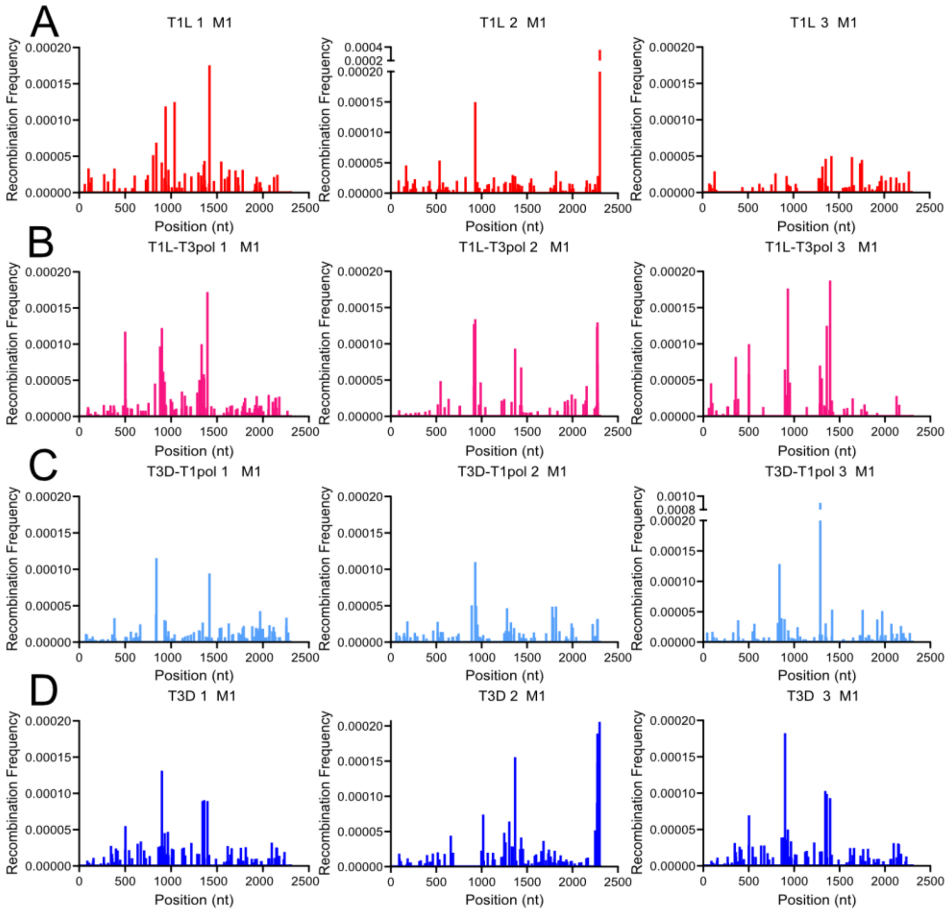


**Figure S16. Reovirus recombination start-site frequency across gene segments: Segment M1.** Recombination start-site frequency at each nucleotide position across the M1 segment from T1L (A), T1L-T3pol (B), T3D-T1pol (C), and T3D (D). Positional recombination frequency of sequenced virion RNA for each individual virus clone is shown.

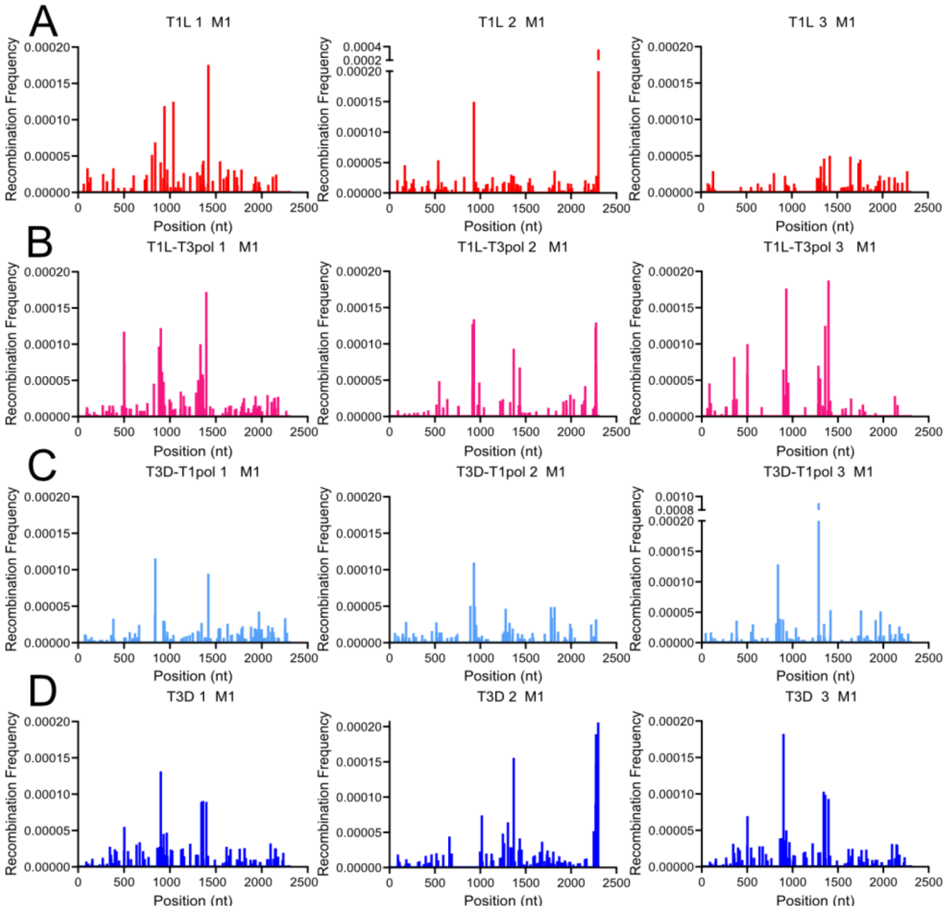


**Figure S17. Reovirus recombination stop-site frequency across gene segments: Segment M1.** Recombination start-site frequency at each nucleotide position across the M1 segment from T1L (A), T1L-T3pol (B), T3D-T1pol (C), and T3D (D). Positional recombination frequency of sequenced virion RNA for each individual virus clone is shown.

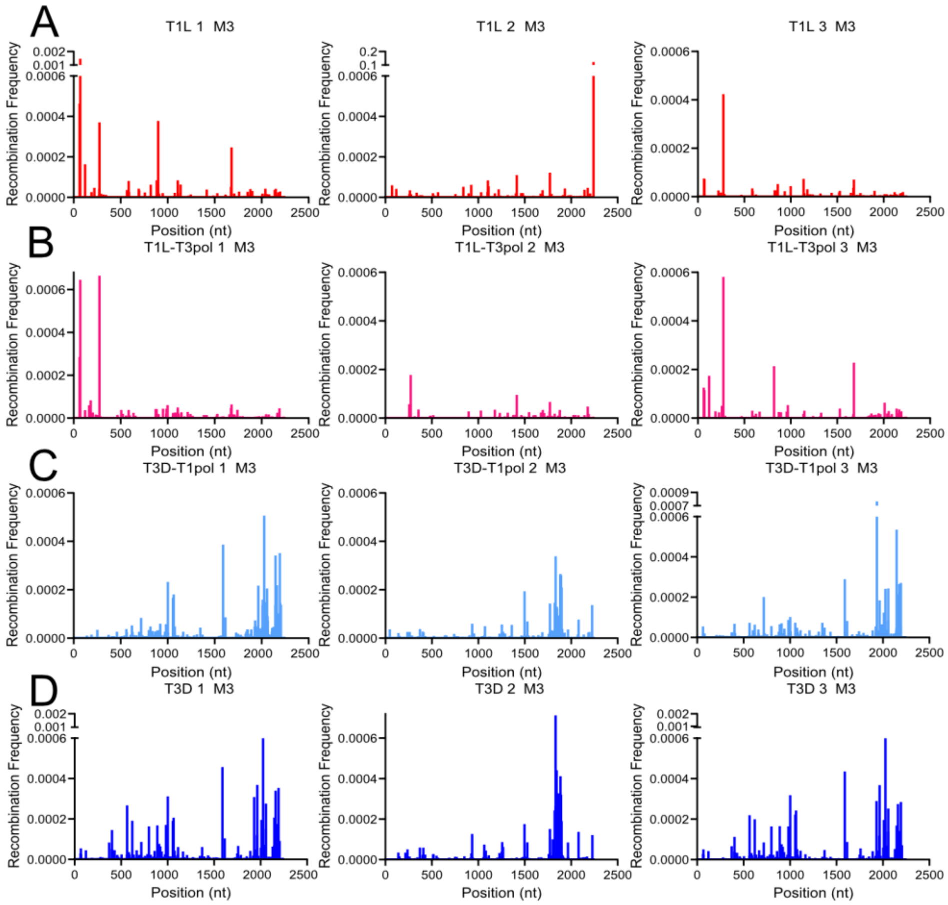


**Figure S18. Reovirus recombination start-site frequency across gene segments: Segment M3.** Recombination start-site frequency at each nucleotide position across the M3 segment from T1L (A), T1L-T3pol (B), T3D-T1pol (C), and T3D (D). Positional recombination frequency of sequenced virion RNA for each individual virus clone is shown.

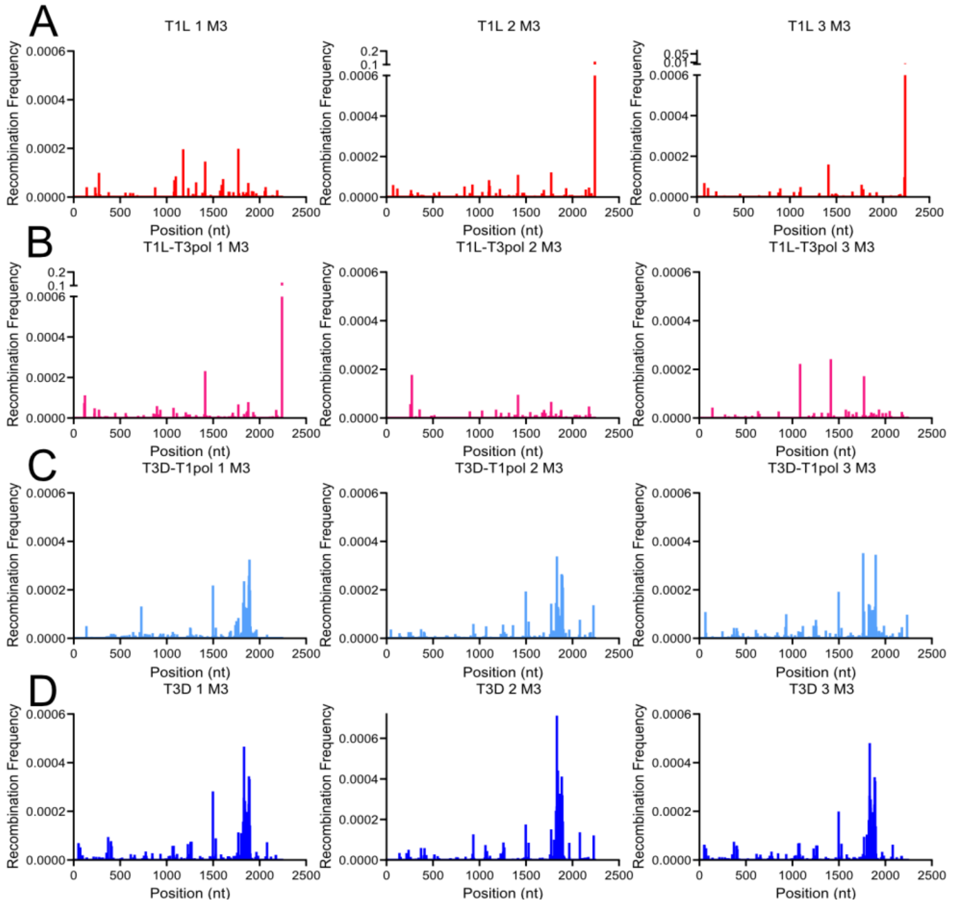


**Figure S19. Reovirus recombination stop-site frequency across gene segments: Segment M3.** Recombination start-site frequency at each nucleotide position across the M3 segment from T1L (A), T1L-T3pol (B), T3D-T1pol (C), and T3D (D). Positional recombination frequency of sequenced virion RNA for each individual virus clone is shown.

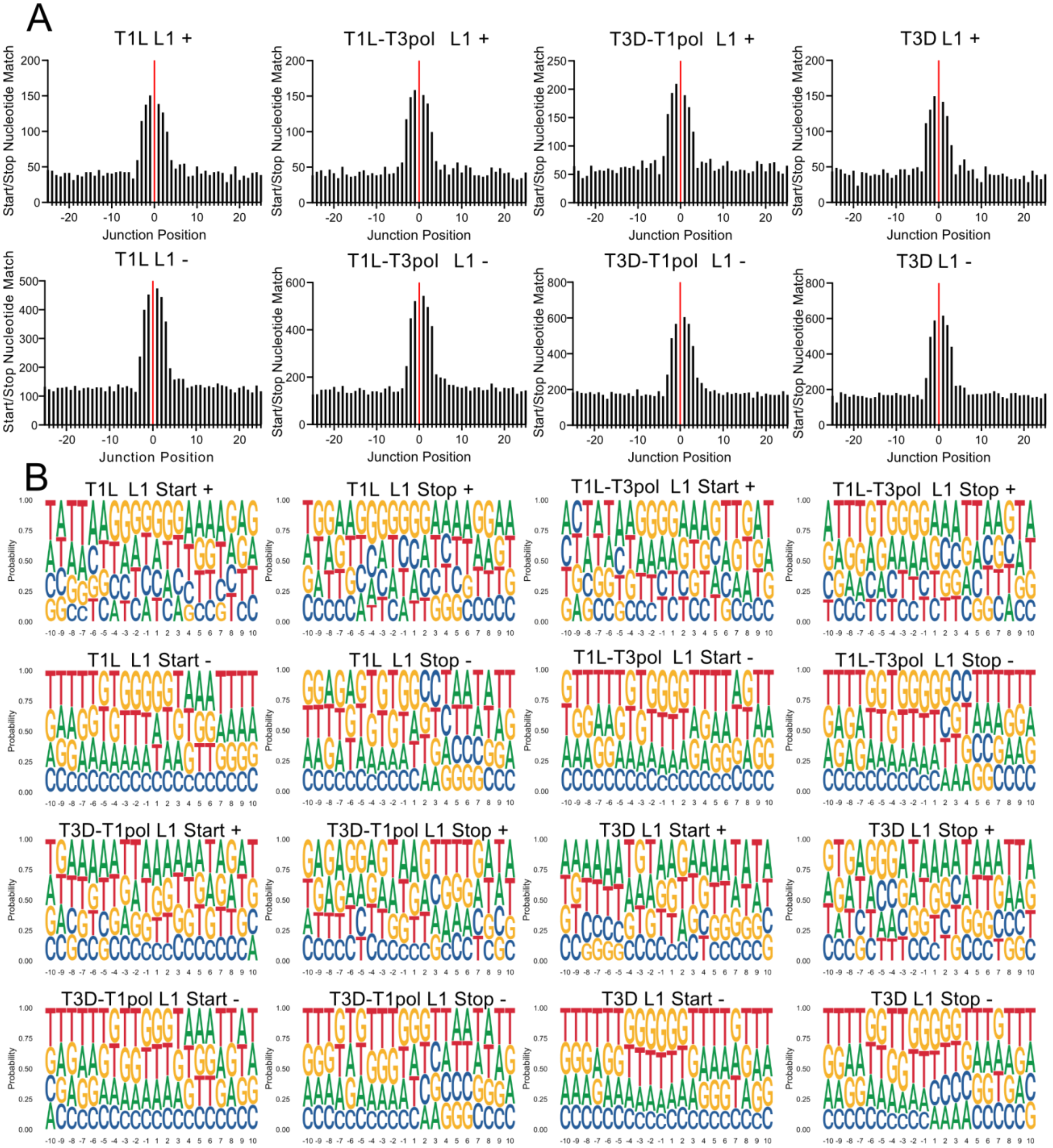

**Figure S20. Microhomology and logo plots for segment L1.** (A) Sequence microhomology for T1L, T1L-T3pol, T3D-T1pol, and T3D. Positionally identical nucleotides in 50-bp regions surrounding start and stop recombination junction sites were quantified for segment L1. (B) Nucleotide composition was calculated as the percent adenosine (A), cytosine (C), guanine (G), and Thymine (T) at each position in a 20-bp region surrounding L1 start and stop recombination junction sites. In (A) and (B), each analyzed virus contains combined data from three independent samples separated by strand, as indicated.

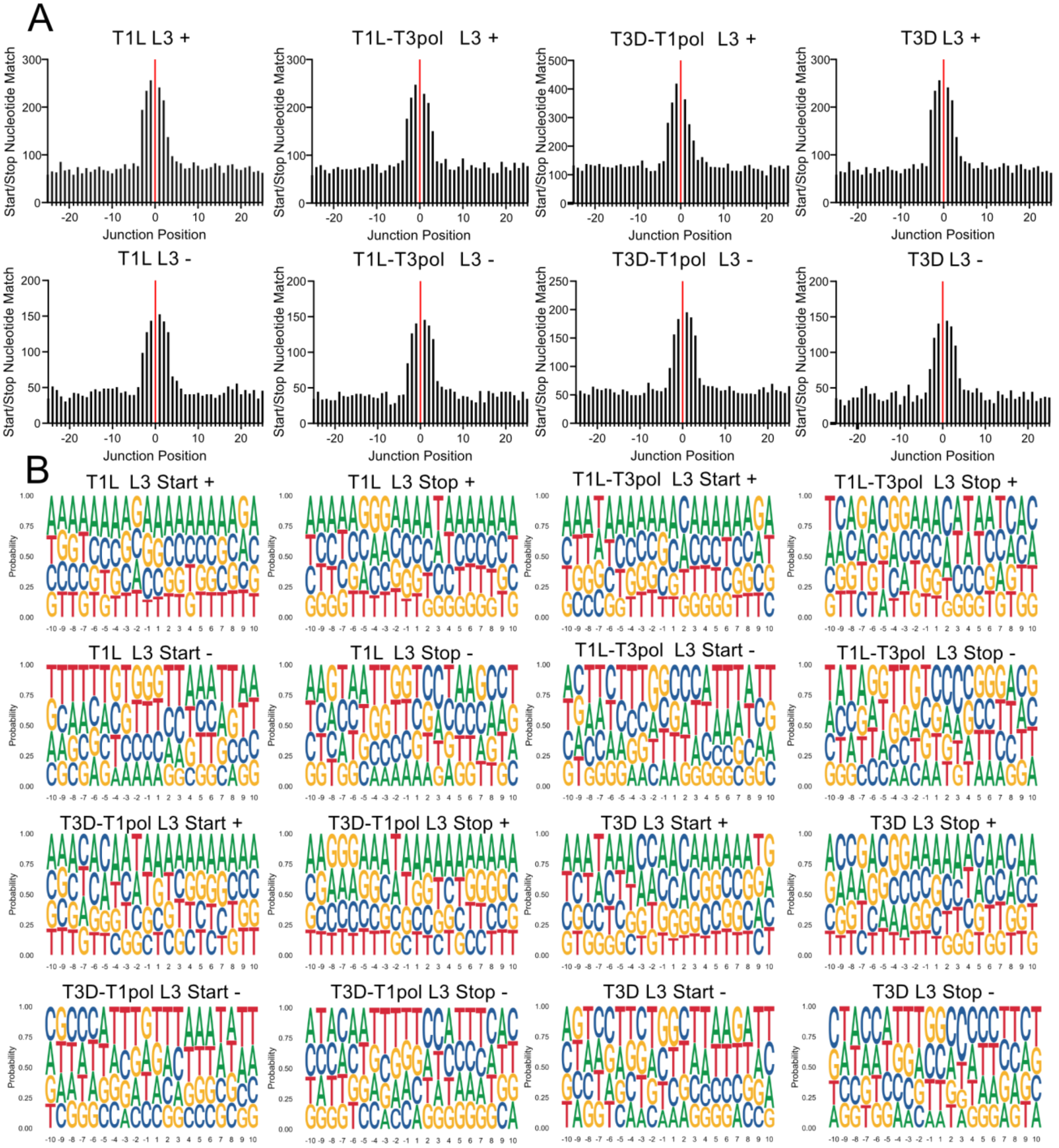


**Figure S21. Microhomology and logo plots for segment L3.** (A) Sequence microhomology for T1L, T1L-T3pol, T3D-T1pol, and T3D. Positionally identical nucleotides in 50-bp regions surrounding start and stop recombination junction sites were quantified for segment L3. (B) Nucleotide composition was calculated as the percent adenosine (A), cytosine (C), guanine (G), and Thymine (T) at each position in a 20-bp region surrounding L3 start and stop recombination junction sites. In (A) and (B), each analyzed virus contains combined data from three independent samples separated by strand, as indicated
